# Screening Lipid Nanoparticles through Structure-Ratio Alignment

**DOI:** 10.64898/2026.07.08.737142

**Authors:** Yoonho Lee, Yunhak Oh, Hoyoung Choi, Chanyoung Park

## Abstract

Lipid Nanoparticles (LNPs) are widely used as delivery systems for nucleic acid therapeutics, where transfection efficiency is determined by both the identities of constituent lipid components and their composition ratios. While prior studies have focused on learning molecular representations for individual components, modeling how multiple components and their ratios jointly influence LNP performance remains underexplored. In this work, we propose STRATA, a framework that models molecule interaction between LNP components, which is known to contribute to LNP transfection efficiency. Our approach is built on two complementary views: (1) a ratio-centric view that captures interaction patterns induced by composition ratios through a transformer with a Ratio-induced Positional Embedding, and (2) a molecule-centric view that incorporates interaction-induced effects into structure-based molecule embeddings. By jointly training and aligning these views, our model integrates molecular structure and composition ratio within a unified framework that captures interaction-driven effects. Experiments demonstrate that our method improves prediction accuracy and generalization to unseen molecules and ratios, highlighting the effectiveness of our approach. Our implementation is available at https://github.com/Yoonho-Lee-AI4Science/STRATA.

## 1 Introduction

Lipid nanoparticles (LNPs) are widely used as delivery systems for nucleic acid cargos such as mRNA, siRNA, and plasmid DNA (pDNA), enabling intracellular delivery while protecting cargos from nuclease-mediated degradation. The success of LNP technology has been demonstrated by its clinical translation, including the first siRNA therapeutic Onpattro[1] and SARS-CoV-2 vaccines such as Spikevax[3] (Moderna) and Comirnaty[19] (Pfizer-BioNTech), which has significantly increased interest in LNP-based delivery systems. A typical LNP formulation, as described in Figure 1 (a), consists of four major components: ionizable lipid (IL), helper lipid (HL), cholesterol (CHOL), and PEG-lipid (PEG). Within an LNP, these components do not act independently but mutually influence [22, 9, 30] one another, a phenomenon we refer to as *molecule interaction*, which collectively gives rise to macroscopic properties such as transfection efficiency. In particular, the composition ratio of the components are known to be associated with their relative spatial arrangement, such as intermolecular distances and positions within the particle, and thereby plays a decisive role in shaping molecule interaction [13, 16]. Consequently, since such interactions are shaped by both the molecular identities of the lipid components and their composition ratios, identifying optimal combinations of these two factors through formulation screening is a key research challenge. However, the underlying mechanisms remain insufficiently understood [5, 15, 12, 17, 24], and the vast combinatorial search space makes experimental screening costly and inefficient. To address these challenges, recent studies have increasingly explored deep learning approaches to predict LNP transfection efficiency and enable high-throughput virtual screening.

**Figure 1:**
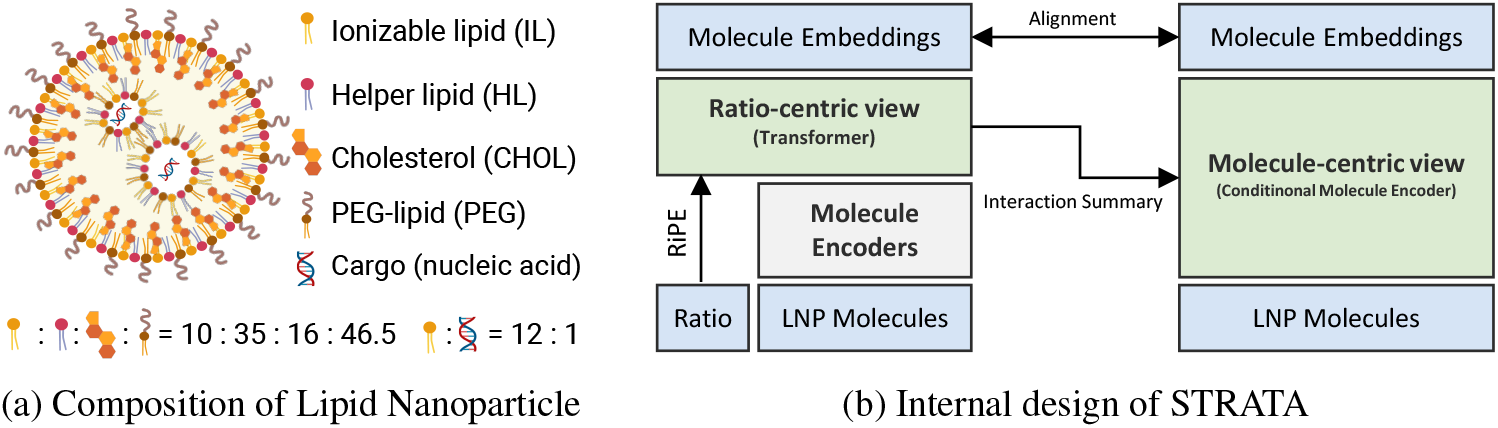
Left figure describes the composition of lipid nanoparticle (LNP), created with BioRender.com. Right figure shows the overall structure of our propose model, STRATA.

Among the four LNP components, ionizable lipids (ILs) have received the most attention due to their critical roles in cargo encapsulation and endosomal escape. Consequently, many deep learning approaches have concentrated on ionizable lipid screening rather than modeling the entire LNP formulation, with a primary focus on how pretrained molecular encoders can be adapted to obtain effective IL representations. Representative strategies include directly fine-tuning pretrained molecular foundation models [27], conducting additional pretraining on lipid-specific datasets [28, 10], and integrating multiple molecular encoders [26].

While substantial progress has been made in learning molecular representations for ionizable lipids, relatively little attention has been paid to how multiple components and their composition ratios should be jointly modeled. In particular, modeling the inter-component interactions induced by a given composition ratio remains underexplored. Existing multi-component approaches treat non-IL components and ratios as auxiliary metadata simply concatenated with IL embeddings [25, 11], or rely on transformer-based aggregation that produces a single summary representation without explicit constraints encouraging interaction modeling [4]. As a result, current approaches struggle to capture the mutual influences among components, and generalize poorly to unseen molecules or composition ratios, an issue further exacerbated by the inherently limited size of LNP datasets.

In this work, we revisit LNP performance prediction through the lens of *molecule interaction*, the mutual influence among components that, as noted above, drives macroscopic outcomes. From this perspective, we propose a novel **ST**ructure-**R**atio **A**ligned **T**ransformer **A**rchitecture, **STRATA**, that models molecule interaction through two complementary views: a ratio-centric view and a molecule-centric view. The ratio-centric view (Figure 1 (b), left tower) focuses on modeling how composition ratios affect interactions among molecules. Specifically, molecule embeddings obtained from a pretrained encoder are updated through a transformer, where we introduce a novel Ratio-induced Positional Embedding (RiPE) to effectively encode ratio information. This design enables the model to reflect interaction patterns driven by composition ratios, producing both updated molecule embeddings and a CLS token that summarizes overall interactions. Building on this interaction summary, the molecule-centric view (Figure 1 (b), right tower) models how interactions influence each molecule with consideration for its molecular structure. To this end, we design a conditional molecule encoder that injects the interaction summary from the ratio-centric view into the encoding process of molecule embeddings from molecular structures. This allows the model to adaptively reflect how interactions influence molecules for a given molecule structure.

These two views complement each other in addressing their respective limitations. The ratio-centric view captures interaction patterns induced by composition ratios, but it operates on fixed molecule embeddings and thus does not adapt the interaction modeling to molecular structure. On the other hand, the molecule-centric view effectively encodes structure dependency through pretrained molecular representations, yet cannot independently model interaction effects arising from composition ratios. By jointly training the two views and aligning their representations, our model enables mutual compensation between them. As a result, both molecular structure and composition ratio are integrated within a unified framework that reflects molecule interaction, leading to improved prediction of LNP performance and enhanced generalization in data-scarce settings.

Our contributions are summarized as follows.

- We propose a new framework that models the process in which LNP components interact under a given composition ratio as the alignment between a ratio-centric view and a molecule-centric view, simultaneously reflecting the effects of molecular structure and composition ratio.
- We design Ratio-induced Positional Embedding (RiPE) so that ratio information can be effectively encoded when reflected through the transformer.
- The proposed model not only achieves outstanding predictive performance, but also demonstrates robust generalization in unseen molecule and unseen ratio settings.

## 2 Related Work

### 2.1 Lipid Nanoparticles and Formulation Screening

Formulation screening for Lipid Nanoparticles (LNPs) aims to predict transfection efficiency given the constituent molecules—ionizable lipid (IL), helper lipid (HL), cholesterol (CHOL), PEG-lipid (PEG), and cargo—and their composition ratios. For nucleic acid cargo, explicit molecular information is typically unavailable and only its ratio is provided, whereas for the other four components, both SMILES representations and composition ratios are given. Only a few studies address LNP formulation screening task. LIFT [11] and LiON [25] adopt metadata-based approaches (Figure 2(a)), where non-IL components are represented via one-hot encodings and composition ratios are aggregated into a vector, which is then concatenated with the output of a molecular encoder for the IL to form an LNP-level representation. However, by treating each piece of information independently in this concatenation scheme, these methods fail to capture the joint effects arising from interactions among components and their composition ratios. COMET [4] employs a UniMol-based [29] molecular encoder to obtain embeddings for the four lipid components, and encodes composition ratios using a Gaussian basis function (GBF) scheme [20] (Figure 2(b)). The ratio embeddings are concatenated with the corresponding molecule embeddings, and a transformer aggregates them into a summary token representing the LNP for downstream prediction. While this approach incorporates both molecular identity and composition ratios, its use of the transformer is limited to information aggregation. As shown in Figure 2(b), only the CLS token is forwarded to the prediction head, and the per-component embeddings updated by the transformer are left unused. Consequently, no training signal directly encourages these embeddings to capture inter-component interactions.

**Figure 2:**
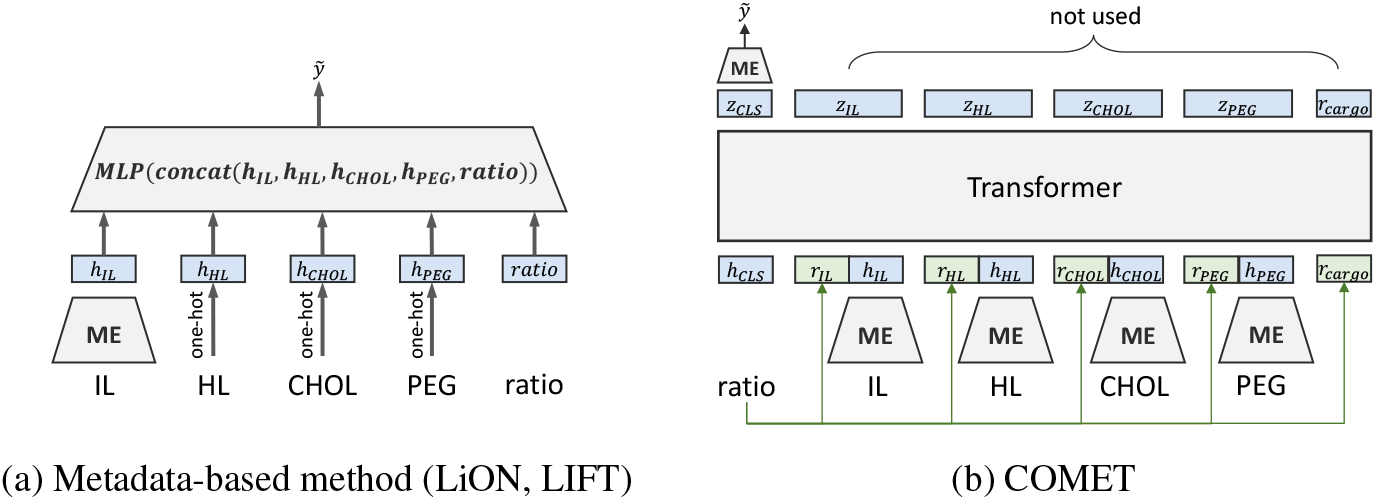
(a) Metadata-based methods append non-IL components (one-hot) and ratios to IL embeddings. (b) COMET aggregates all components via a transformer but does not use updated molecule embeddings, lacking constraints for interaction modeling. ME denotes molecule encoders.

### 2.2 Positional Encoding

Positional encoding has been studied extensively in the sequence-modeling literature, and the methods developed there provide natural starting points for encoding compositional ratios in LNP formulations. The original Transformer [23] adopts absolute positional encoding (APE) using sinusoidal functions of the absolute index, which injects a unique vector for every position. COMET [4] instead encodes each ratio through a Gaussian basis function (GBF) [20] scheme: a scalar ratio *r* ∈ R is passed through *d* radial basis functions parameterized by learnable means 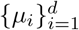 and standard deviations 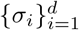, yielding a *d*-dimensional embedding

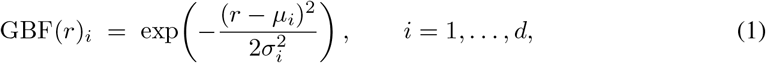

which is then concatenated to the corresponding molecular embedding before being fed into the model. In transformer-based interaction modeling, attention mechanisms aim to reflect the influence between pairs of molecules. Since the ratio between two molecules is a relative property, APE and GBF, which rely on absolute encoding schemes, are limited in capturing the relative nature of ratios required for interaction modeling via attention.

Rotary Position Embedding (RoPE) [21] addresses this limitation by constructing position-dependent transformations *f*_*q*_(*x*_*m*_, *m*) and *f*_*k*_(*x*_*n*_, *n*) such that the query–key inner product depends on positions only through their relative offset:

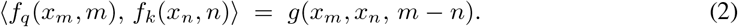

Concretely, RoPE realizes this property by applying a rotation matrix parameterized by *m* − *n*, so that any uniform translation of the indices leaves the inner product invariant. While this elegantly captures relativeness for token positions, it does not match the intrinsic character of compositional ratios. The defining property of a ratio is *multiplicative* relativeness (*m/n*) rather than *additive* relativeness (*m* − *n*): the pairs 10 : 20 and 20 : 30 share the same additive offset and would therefore yield identical relative encodings under RoPE, yet they correspond to fundamentally different compositions (1 : 2 versus 2 : 3). In short, none of APE, GBF, or RoPE encodes the inductive bias that compositional ratios call for—namely, that the meaning of a ratio is preserved under multiplicative rescaling rather than additive translation. This mismatch between the intrinsic nature of ratios and the inductive bias of existing positional encodings motivates the ratio-aware encoding scheme we develop in the next section.

## 3 Method

STRATA comprises three components, illustrated in Figure 3. The *ratio-centric view* (Figure 3 (a)) performs interaction modeling through an interaction transformer equipped with a Ratio-induced Positional Embedding (RiPE), which encodes composition ratios as an inductive bias and yields an interaction summary over the formulation. The *molecule-centric view* (Figure 3 (b)) then re-derives each lipid’s embedding from its molecular structure using a Conditional Molecule Encoder (CME; Figure 3 (d)) that takes the interaction summary as a conditioning signal, so that structural information is updated under the local environment shaped by interactions. Because the two views describe the same interaction phenomenon from complementary perspectives, we further introduce an *alignment objective* (Figure 3 (c)) that regularizes the per-component representations of the two views to remain consistent, allowing them to compensate for each other’s limitations.

**Figure 3:**
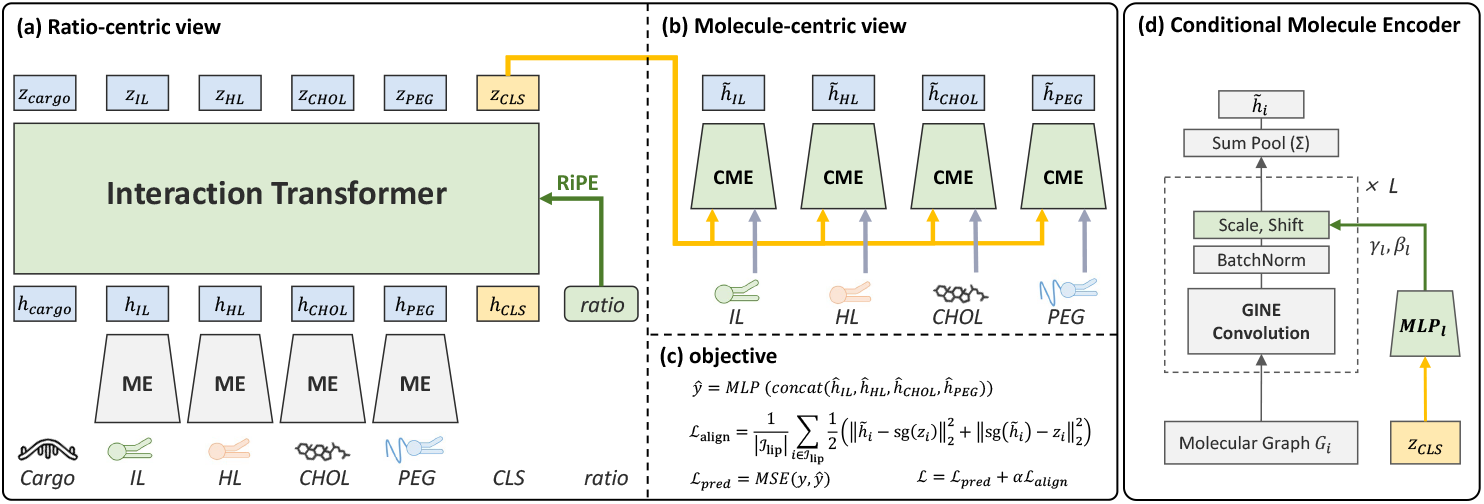
Overall description of STRATA. Interaction summary (shown in yellow) from the ratio-centric view (a) serves as the condition for the molecule-centric view (b). The Conditional Molecule Encoder (d) produces condition-dependent molecule embeddings, and alignment (c) bridges the two views to compensate for each other’s limitations. ME and CME denote the Molecule Encoder and Conditional Molecule Encoder, respectively. Our core design components are highlighted in green.

### Notation

We denote an LNP formulation by the tuple 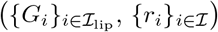, where *I* = {IL, HL, CHOL, PEG, cargo} is the set of formulation components and *I*_lip_ = {IL, HL, CHOL, PEG} ⊂ *I* is the subset of lipid components for which molecular structures are available. For each lipid *i* ∈ *I*_lip_, the molecule is represented as a graph *G*_*i*_ = (*V*_*i*_, *E*_*i*_) with atom set *V*_*i*_ and bond set *E*_*i*_; the cargo is treated as structure-less and enters the model only through its ratio *r*_cargo_. Each component *i* ∈ *I* is associated with a positive compositional ratio *r*_*i*_ ∈ R_*>*0_. The scalar regression target *y* ∈ R denotes the transfection efficiency, and *ŷ* denotes the model prediction of *y*. Throughout the paper, *d* denotes the hidden dimension shared by all embeddings.

#### 3.1 Ratio-centric modeling with interaction transformer

The interaction transformer (Figure 3 (a)) takes as input a CLS token, the four lipid component tokens, the cargo token, and their associated compositional ratios. The CLS token serves as a learnable summary slot whose output we treat as the formulation-level interaction summary. The full input sequence is *X*^(0)^ =[*h*_cargo_, *h*_IL_, *h*_HL_, *h*_CHOL_, *h*_PEG_, *h*_CLS_] ∈ R^6*×d*^.

For the four lipid component token embeddings, we adopt the pretrained molecule encoder of AG-ILE [28], a GINE-based [14] pretrained graph neural network, and keep it frozen throughout training. The cargo and CLS tokens are learnable embeddings. Because our dataset provides no per-formulation information about the cargo, the same cargo token *h*_cargo_ is shared across all samples. We nonetheless retain it as an explicit token so that its compositional ratio is injected through the positional encoding and its interactions with the lipid tokens are captured via attention. From the interaction transformer, which incorporates molecule token embeddings derived from the AGILE molecular encoder, along with learnable cargo, CLS embeddings and RiPE, we obtain updated molecule embeddings *z*_IL_, *z*_HL_, *z*_CHOL_, *z*_PEG_, *z*_cargo_ ∈ R^*d*^ and an interaction-level summary representation *z*_CLS_ ∈ R^*d*^.

To let the attention between two component tokens reflect the relative effect of their compositional ratios, we design RiPE as a variant of RoPE that adjusts attention scores through ratio-dependent rotation matrices. As discussed in Section 2.2, standard RoPE computes its rotation from the *difference* of two positions and is therefore not aligned with the multiplicative nature of ratios. We resolve this with a single modification: before forming the rotation matrix, we apply a logarithmic transformation to the ratio that defines each token’s position. Because the logarithm turns addition/subtraction into multiplication/division, this simple change is sufficient to align the rotation geometry with the intrinsic nature of ratios.

##### Formal definition of RiPE

Let *W*_*q*_, *W*_*k*_ ∈ R^*d×d*^ be the query and key projection matrices, and let Θ = (*θ*_1_, …, *θ*_*d/*2_) with *θ*_*t*_ = 10000^*−*2(*t−*1)*/d*^ be the RoPE frequency schedule. For a scalar *p* ∈ R, define the block-diagonal rotation operator

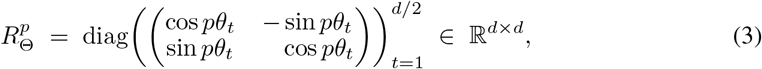

which rotates each consecutive pair of feature dimensions by an angle proportional to *p*. Standard RoPE sets *p* = *m* for an integer position *m*. RiPE instead sets *p* = log *m* where *m* ∈ R_*>*0_ is a compositional ratio: 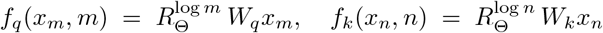.

###### Proposition 1

(Properties of RiPE). *The proposed Ratio-induced Positional Embedding (RiPE) satisfies the following three properties:*

1. ***Ratio dependence***. *The attention score induced by RiPE depends on positions m, n through the ratio m/n: f*_*q*_(*x*_*m*_, *m*), *f*_*k*_(*x*_*n*_, *n*) = *g*(*x*_*m*_, *x*_*n*_, *m/n*).
2. ***Scale invariance***. *For all c >* 0, *the attention score is invariant under the joint rescaling* (*m, n*) ↦ (*cm, cn*). *Equivalently, per-token RiPE encoding transforms equivariantly under the multiplicative group* (R_*>*0_, ×), *with resulting per-token rotations canceling in the inner product*.
3. ***Compositionality of ratios***. *The pairwise rotations induced by RiPE compose multiplicatively along any chain of positions a* → *b* → *c, recovering the direct ratio c/a from c/b and b/a*.

Although the arguments are simple and intuitive, we provide short proofs in Appendix A to make explicit that RiPE reflects the nature of ratios. Despite its simplicity, this ratio-aware design translates into substantial empirical gains on unseen ratios, as demonstrated in Section 4. Further discussion on RiPE is provided in Appendix A.6.

#### 3.2 Molecule-centric modeling with Conditional Molecule Encoder (CME)

Updating molecule embeddings through molecule interaction requires accounting for not only compositional ratios but also each component’s own molecular structure. Because the ratio-centric view does not let a token’s embedding be revisited in light of its own structure during the update, the goal of the molecule-centric view is to perform interaction modeling that explicitly accounts for molecular structure. The interaction transformer’s *z*_CLS_ already summarizes the surrounding influence on each component, so we treat it as a conditioning signal and construct a *conditional molecule encoder* (CME) that, given a molecule’s local environment as condition, encodes the molecule’s properties from its structure. We reuse the AGILE encoder introduced earlier, but make it learnable in this stage, and enable layer-wise conditional modulation through the FiLM [18] structure. Note that the CME and the molecule encoder in Section 3.1 share the same parameter initialization but do not share weights.

For formal description, let 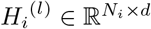 denote node features at layer *l* of the encoder, where *N*_*i*_ is the number of atoms in *G*_*i*_. A standard GINE layer applies message passing followed by batch normalization and a ReLU; we follow the FiLM [18] recipe and insert an modulation between the normalization and the nonlinearity (Figure 3 (d)):

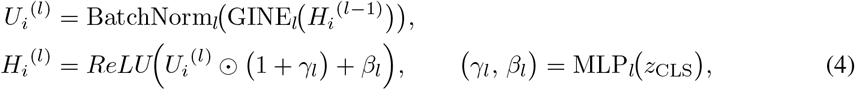

where ⊙ is the elementwise product, *γ*_*l*_, *β*_*l*_ ∈ R^*d*^ are produced by a small layer-specific MLP applied to the summary *z*_CLS_. After *L* blocks, node features are sum-pooled to obtain the graph-level embedding 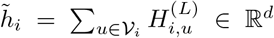 where 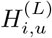 denotes embedding of a node *u* ∈ *G*_*i*_.

#### 3.3 Training Objective

For transfection efficiency *y* prediction, we concatenate the embeddings of the four lipid components obtained from the conditional molecule encoder: 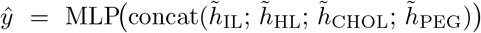.

We optimize the model using a mean squared error loss L_pred_ = MSE(*y, ŷ*). Because the two views describe the same molecular interactions from complementary perspectives, we regularize 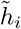 not to deviate far from *z*_*i*_. Since *z*_*i*_ already encodes formulation-level interaction context through attention, this regularization encourages the conditional molecule encoder to actively utilize the interaction summary *z*_CLS_ it receives as conditioning, rather than relying solely on the molecular graph. Conversely, since 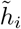 is derived directly from the molecular graph, aligning toward it encourages the interaction transformer to produce per-component representations that remain grounded in the underlying molecular structure. This mutual regularization allows the two views to compensate for each other’s limitations.

Thus, we introduce a symmetric stop-gradient alignment loss 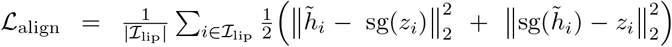, where sg(·) denotes the stop-gradient operator. Since both views are learnable, the stop-gradient stabilizes training by decoupling the updates: in each term one view serves as a fixed target while the other optimizes toward it, so that each branch is guided to reflect the information already captured by its counterpart. The final objective is L = L_pred_ + *α* L_align_.

## 4 Experiment

We conduct experiments to answer the following research questions:

- **RQ1**. Does STRATA perform well on transfection efficiency prediction tasks, particularly for unseen molecules or composition ratios? (§4.2)
- **RQ2**. Does our proposed Ratio-induced Positional Encoding (RiPE) contribute to performance gains over alternative positional encoding schemes? (§4.3)
- **RQ3**. Do STRATA’s predictions faithfully reflect formulation ratios? In other words, are its predictions sensitive to variations in the ratios? (§4.4)
- **RQ4**. To what extent does STRATA capture transferable representations of LNP formulations across cell types? (§4.5)

### 4.1 Experimental Setup

#### Datasets

We evaluate our model on four transfection efficiency datasets: A549, YZ2022, LC2024 (N2a), and LC2024 (HEK293), which span diverse cell lines and experimental conditions. The A549 dataset was obtained from [25], YZ2022 from [8], and LC2024 (N2a) and LC2024 (HEK293) from [6]. Preprocessing details and dataset statistics for each dataset are provided in Appendix B.1.

#### Evaluation splits

To rigorously assess generalization, we evaluate under seven splits. The *random split* serves as the in-distribution setting. The *unseen molecule split* is constructed such that there is no overlap of molecules across the train/validation/test sets. Specifically, since each dataset focuses on different molecule types, we use unseen ionizable lipids for A549 and unseen helper lipids for the remaining datasets. The five *unseen ratio splits* (IL, HL, CHOL, PEG, and IL-to-cargo) hold out specific ranges of each formulation ratio so that test-time ratio combinations are unseen during training, directly probing compositional generalization. For the unseen IL-to-cargo split, only a limited number of distinct values (3–4) are available; therefore, we adopt a 1:1:1 train/validation/test split. For all other split settings, the data are divided into train/validation/test with a ratio of 3:1:1.

#### Metrics

Following prior work, we report mean squared error (MSE; lower is better) and Pearson correlation coefficient (PCC; higher is better). We evaluate each model over 10 different random seeds and report the mean performance along with the standard deviation. Model performance is reported using the parameters that achieve the lowest MSE on the validation set.

#### Baselines

Our study focuses on how to integrate component identities and their ratios rather than on developing a new molecular encoder. To isolate the effect of this integration strategy from confounding factors such as the choice of molecular encoder and the amount of lipid-specific pretraining, we construct AGILE-based variants of all baselines. AGILE-Metadata represents the approach used in LiON [25] and LIFT [11] (Figure 2 (a)). AGILE-MLP is a simple variant derived from AGILE-Metadata for experimental purposes; it processes ratio information in the same way as AGILE-Metadata but differs in that all components are passed through the molecule encoder. While AGILE-MLP still treats ratio as auxiliary information and does not explicitly model interactions, it can be viewed as a variant that treats all four components as equally important as the ionizable lipid. AGILE-COMET is a variant of COMET [4] (Figure 2 (b)) in which the original molecule encoder is replaced with AGILE. Implementation details and hyperparameter search spaces are available at Appendix B.2.

### 4.2 Results for Transfection Efficiency Prediction (RQ1)

Table 1 reports performance across all splits and datasets. We compute the relative performance gain of STRATA (*p*_ours_) over the best-performing baseline (*p*_base_) as |*p*_base_ − *p*_ours_|*/p*_base_ × 100 (%). We assign a positive value when our model outperforms the best baseline, and a negative value when the best baseline achieves better performance. We have the following observations: **1)** The proposed STRATA achieves the best performance on almost all datasets and split settings for both metrics. This indicates that STRATA exhibits stronger generalization than competing models, effectively handling unseen molecules and ratio combinations encountered at test time. **2)** Although COMET adopts a transformer-based architecture similar to STRATA, it performs worse than AGILE-MLP, which uses a simple MLP structure. This suggests that naively relying on a transformer alone is insufficient. In contrast, our model effectively addresses this limitation by aligning two complementary views, which implicitly model interactions from different perspectives.

**Table 1:**
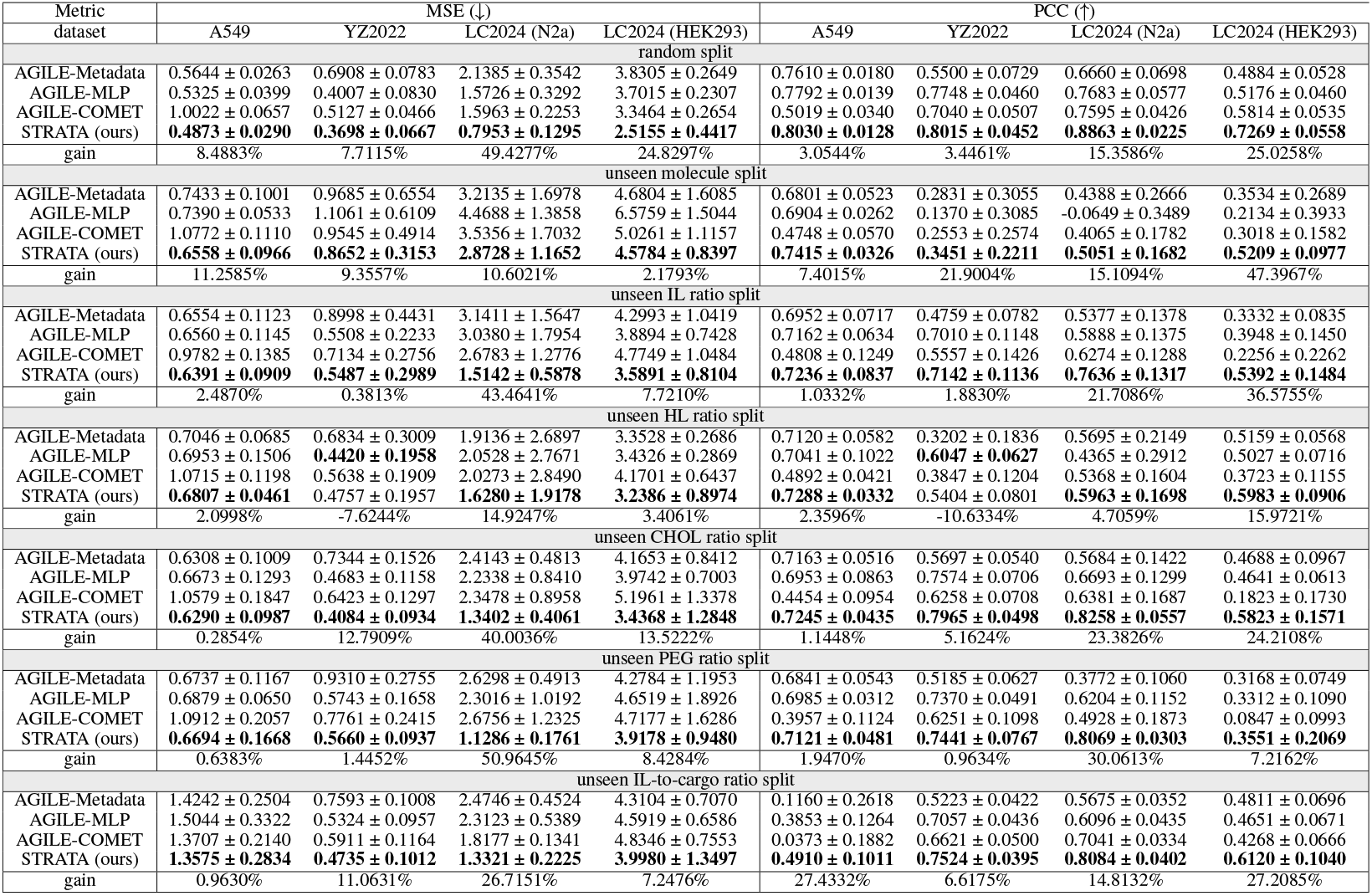
Model performance on transfection efficiency prediction task. The best performance for each metric is highlighted in bold.

### 4.3 Evaluation of RiPE’s Effectiveness (RQ2)

To verify the effectiveness of Ratio-induced Positional Embedding (RiPE), we evaluated STRATA while replacing its positional encoding scheme with absolute positional encoding [23] (APE), Rotary Position Embedding [21] (RoPE), and the Gaussian basis function (GBF) used in COMET. In addition, to isolate the effect of using the ratio itself as a positional signal, we constructed a variant in which a learnable embedding is assigned to each component type (IL, HL, CHOL, PEG) and added to the corresponding token instead of using ratios; this variant is denoted as ‘random’ in the table. The results are summarized in Table 2. We make the following observations: **1)** Regardless of the split or dataset, RiPE consistently outperforms the other positional encoding schemes. Comparing these results with Table 1 reveals that RiPE plays a more critical role than one might expect: when RiPE is replaced with alternative positional encoding schemes, STRATA often fails to surpass even simple baselines such as AGILE-MLP. This indicates that positional encoding contributes substantially to overall performance, and that an appropriate modeling strategy tailored to ratios is essential. **2)** Despite the similarity between RoPE and RiPE, they exhibit a surprisingly large performance gap. RoPE is designed to capture the relative relationship between two tokens, but it is built on distance (*m* − *n*), which does not align with the multiplicative nature (*m/n*) of ratios. This highlights that faithfully reflecting the inherent nature of ratios is critically important. Our RiPE, despite its simplicity, is effective precisely because it embeds an inductive bias that matches the multiplicative nature of ratios. The full table, including additional splits such as unseen HL, CHOL, PEG, and IL-to-cargo splits, is provided in Table 10 of Appendix C.4.

**Table 2:** The performance of STRATA with different positional encodings.

| Metric<br>dataset | MSE (↓) | | | | PCC (↑) | | | |
| --- | --- | --- | --- | --- | --- | --- | --- | --- |
|  | A549 | YZ2022 | LC2024 (N2a) | LC2024 (HEK293) | A549 | YZ2022 | LC2024 (N2a) | LC2024 (HEK293) |
| random split |  |  |  |  |  |  |  |  |
| random | 0.7197 ± 0.0424 | 0.8647 ± 0.1455 | 3.4426 ± 0.4141 | 4.7942 ± 0.2014 | 0.6796 ± 0.0282 | 0.3670 ± 0.0495 | 0.2886 ± 0.0646 | 0.2428 ± 0.0384 |
| APE | 0.5503 ± 0.0200 | 0.4624 ± 0.0620 | 1.1075 ± 0.2099 | 2.6291 ± 0.5830 | 0.7767 ± 0.0157 | 0.7469 ± 0.0587 | 0.8487 ± 0.0289 | 0.7132 ± 0.0642 |
| RoPE | 0.5511 ± 0.0420 | 0.4927 ± 0.0983 | 1.5120 ± 0.2974 | 2.9154 ± 0.4961 | 0.7711 ± 0.0232 | 0.7290 ± 0.0576 | 0.7942 ± 0.0462 | 0.6892 ± 0.0606 |
| GBF (comet) | 0.5778 ± 0.0279 | 0.4817 ± 0.1165 | 1.1939 ± 0.1555 | 2.8338 ± 0.4584 | 0.7606 ± 0.0189 | 0.7280 ± 0.0703 | 0.8325 ± 0.0319 | 0.6832 ± 0.0531 |
| RiPE (ours) | 0.4873 ± 0.0290 | 0.3698 ± 0.0667 | 0.7953 ± 0.1295 | 2.5155 ± 0.4417 | 0.8030 ± 0.0128 | 0.8015 ± 0.0452 | 0.8863 ± 0.0225 | 0.7269 ± 0.0558 |
| gain | 11.4483% | 20.0260% | 28.1896% | 4.3209% | 3.3861% | 7.3102% | 4.4303% | 1.9209% |
| unseen molecule split |  |  |  |  |  |  |  |  |
| random | 1.0007 ± 0.0877 | 1.0731 ± 0.6086 | 3.7470 ± 1.2825 | 6.7951 ± 1.3772 | 0.5504 ± 0.0576 | -0.0538 ± 0.1673 | -0.0418 ± 0.1014 | 0.0593 ± 0.1040 |
| APE | 0.6947 ± 0.1012 | 0.9871 ± 0.4978 | 2.9962 ± 1.7400 | 5.5448 ± 2.2166 | 0.7292 ± 0.0241 | 0.2741 ± 0.1713 | 0.4863 ± 0.1461 | 0.4115 ± 0.2084 |
| RoPE | 0.6917 ± 0.0866 | 1.0367 ± 0.5597 | 3.8003 ± 1.5614 | 5.1754 ± 1.5710 | 0.7252 ± 0.0248 | 0.2559 ± 0.2267 | 0.4153 ± 0.2369 | 0.4553 ± 0.1534 |
| GBF (comet) | 0.7543 ± 0.1319 | 1.0688 ± 0.5353 | 3.2946 ± 1.5187 | 6.3435 ± 1.7618 | 0.6977 ± 0.0369 | 0.0543 ± 0.1936 | 0.3902 ± 0.2189 | 0.3409 ± 0.1429 |
| RiPE (ours) | 0.6558 ± 0.0966 | 0.8652 ± 0.3153 | 2.8728 ± 1.1652 | 4.5784 ± 0.8397 | 0.7415 ± 0.0326 | 0.3451 ± 0.2211 | 0.5051 ± 0.1682 | 0.5209 ± 0.0977 |
| gain | 5.1901% | 12.3493% | 4.1186% | 11.5353% | 1.6868% | 25.9030% | 3.8659% | 14.4081% |
| unseen IL ratio split |  |  |  |  |  |  |  |  |
| random | 0.6539 ± 0.0573 | 1.1240 ± 0.5371 | 4.2737 ± 1.4860 | 5.0135 ± 1.0587 | 0.7005 ± 0.0305 | 0.3564 ± 0.1680 | 0.2666 ± 0.2224 | 0.1744 ± 0.2426 |
| APE | 0.7762 ± 0.1465 | 0.9678 ± 0.4459 | 2.6219 ± 1.6446 | 5.3475 ± 2.4358 | 0.6475 ± 0.0868 | 0.4825 ± 0.1294 | 0.6469 ± 0.1393 | 0.3988 ± 0.1498 |
| RoPE | 0.9527 ± 0.1718 | 0.9229 ± 0.3500 | 2.3285 ± 0.7900 | 4.6465 ± 1.0279 | 0.5189 ± 0.1318 | 0.5362 ± 0.1622 | 0.6430 ± 0.1645 | 0.3561 ± 0.1288 |
| GBF (comet) | 0.7902 ± 0.1710 | 1.0093 ± 0.4835 | 2.6038 ± 1.2074 | 4.3791 ± 1.1059 | 0.6416 ± 0.0836 | 0.4757 ± 0.1578 | 0.6389 ± 0.1729 | 0.4236 ± 0.1050 |
| RiPE (ours) | 0.6391 ± 0.0909 | 0.5487 ± 0.2989 | 1.5142 ± 0.5878 | 3.5891 ± 0.8104 | 0.7236 ± 0.0837 | 0.7142 ± 0.1136 | 0.7636 ± 0.1317 | 0.5392 ± 0.1484 |
| gain | 2.2633% | 40.5461% | 34.9710% | 18.0402% | 3.2976% | 33.1966% | 18.0399% | 27.2899% |

### 4.4 Probing Ratio Sensitivity in Predictions (RQ3)

Effective formulation screening requires the model to faithfully capture how transfection efficiency changes with formulation ratios. We therefore investigate how sensitively our model’s predictions respond to differences in formulation ratios. To examine this property, we measure *ratio sensitivity*: for pairs of formulations *p, q* sharing the same molecular backbone, we compute the magnitude of their ratio difference as 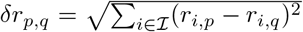, where *r*_*i,p*_ and *r*_*i,q*_ denote the molar ratios of formulation component *i* ∈ *I* in samples *p* and *q*, respectively. We then bin sample pairs by *δr*_*p,q*_ (0–20%, 20–40%, …, 80–100%) and report how closely the predicted efficiency change tracks the ground-truth change within each bin, measured by mean absolute error MAE(*δ*_*y*_, *δ*_*ŷ*_), where *δ*_*y*_ = *y*_*p*_ − *y*_*q*_ and *δ*_*ŷ*_ = *ŷ*_*p*_ − *ŷ*_*q*_ denote the label and prediction differences, respectively.

Figure 4 (top) compares STRATA with baseline architectures across the random, unseen IL ratio, and unseen CHOL ratio settings on the LC2024 (N2a) dataset. STRATA consistently outperforms the baselines and maintains a stable MAE even as the ratio difference grows, indicating that its predictions remain sensitive to ratio variations across the entire range. In contrast, the baselines exhibit substantially degraded sensitivity, suggesting that they largely fail to track ratio-induced changes. Figure 4 (bottom) reports the results of replacing the positional encoding within STRATA. While most variants achieve comparable ratio sensitivity under the random split, only RiPE preserves this sensitivity under the unseen ratio settings, whereas the alternatives deteriorate sharply. This further confirms that our proposed RiPE is critical for faithfully capturing ratio information in formulation-level prediction.

**Figure 4:**
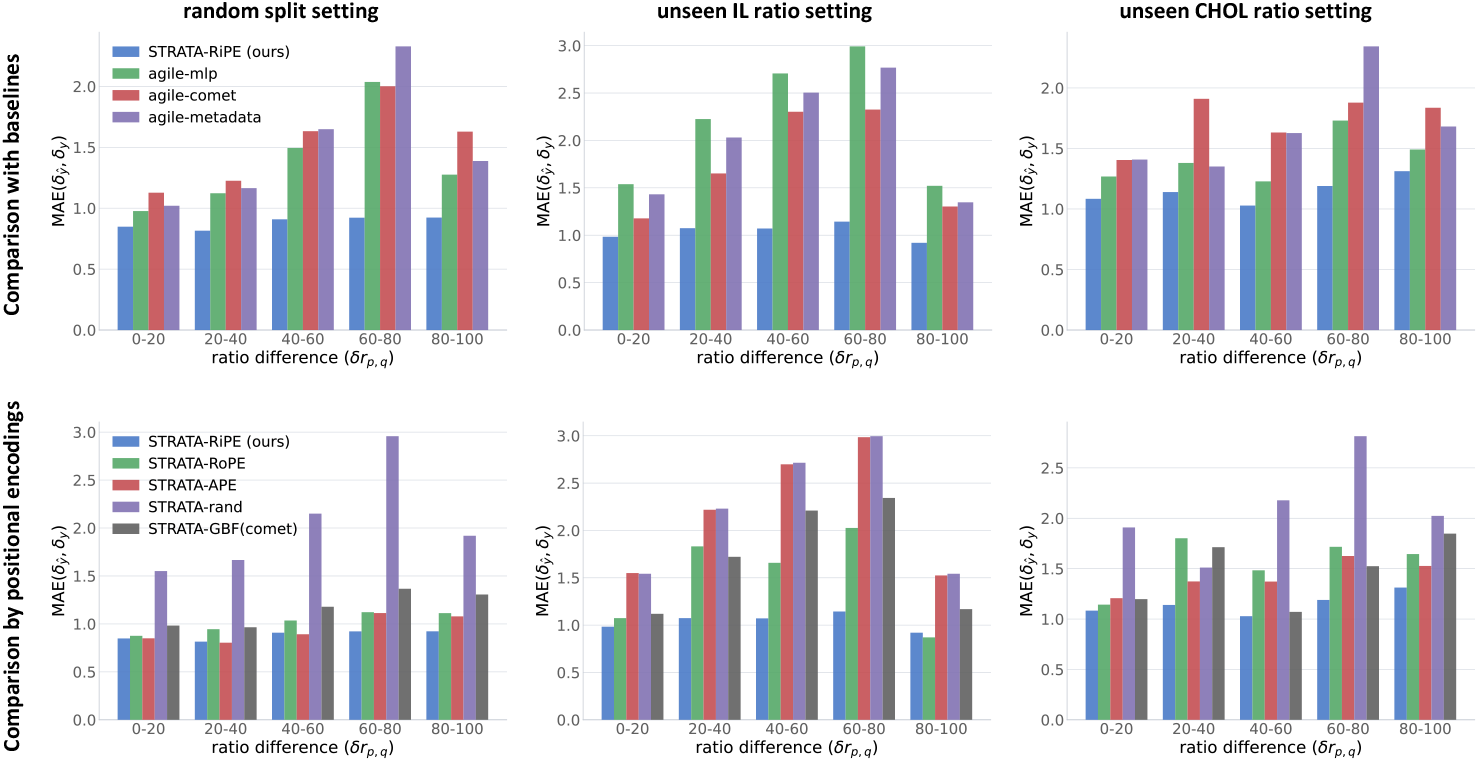
Ratio sensitivity analysis. The first row compares STRATA against baselines, while the second row compares different positional encoding schemes within STRATA. Our model, STRATA, is colored in **blue.**Legends were displayed only on the left figure of each row.

Figure 5 provides a qualitative analysis via t-SNE visualizations of LNP embeddings, colored by the ratio of each formulation component. We visualize the embeddings fed directly into the prediction MLP: for STRATA, we concatenate the four output representations from the CME module, while for AGILE-COMET we use the CLS token representation, following its original design. Since LNP embeddings are shaped by the joint influence of multiple factors, perfect ratio-aligned clustering is not expected; nonetheless, STRATA’s embeddings exhibit clear local clustering structure with respect to each ratio component, whereas AGILE-COMET’s embeddings show only weak clustering or no discernible clustering at all. To quantify how strongly similar ratio values cluster in the embedding, we annotate each t-SNE plot (upper-right corner) with Moran’s I [2, 7], a standard measure of spatial autocorrelation ranging from −1 (dispersed) through 0 (random) to +1 (perfectly clustered). It is computed on a row-normalized kNN graph (*k* = 10) built in the t-SNE space, and captures the degree to which neighboring points share similar values. The Moran’s I values are consistently higher for STRATA than for AGILE-COMET across all five ratios, indicating stronger local coherence of compositional similarity in the STRATA embedding. This qualitative evidence further corroborates that STRATA effectively internalizes the influence of formulation ratios.

**Figure 5:**
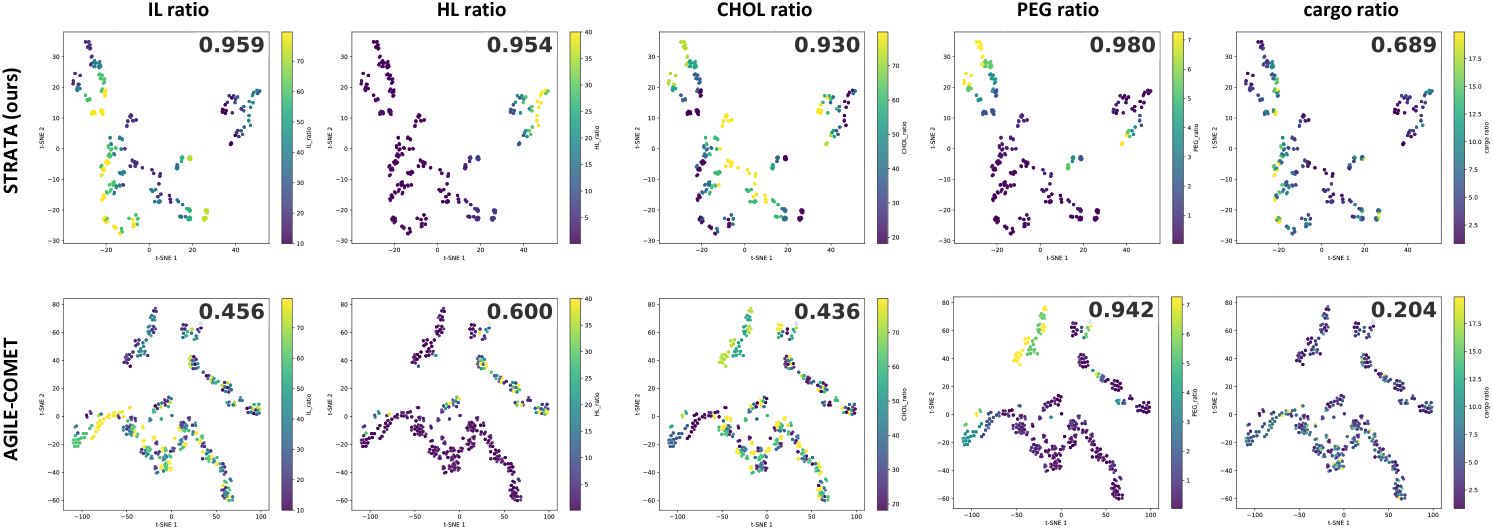
t-SNE visualization of LNP embeddings. The first row shows results from our model, STRATA, and the second row shows results from AGILE-COMET. Each column corresponds to results labeled by five different ratios. The value represents Moran’s I.

### 4.5 Cross-cell type transferability (RQ4)

If the interaction modeling in STRATA captures intrinsic properties of LNP formulations rather than overfitting to cell-type specific patterns in the training data, the learned representations should remain informative for predicting transfection efficiency across different cell types. To test this, we pretrain models on LC2024 (HEK293), freeze all parameters except the prediction head, and fine-tune on four unseen cell types with random split setting. Table 3 summarizes the transfer performance. STRATA consistently outperforms all baselines by a substantial margin across four unseen cell types (ARPE19, B16, N2a, PC3) in both MSE and PCC, demonstrating that our model learns more transferable representations of LNP formulations. However, comparing the transfer results on LC2024 (N2a) with STRATA trained with all parameters learnable in Table 1 reveals a non-negligible performance gap, suggesting that cell-type specific factors may also play a role that is not fully captured by formulation-level representations alone.

**Table 3:** Cross-cell type transfer performance trained on LC2024 (HEK293) and finetuning only prediction heads.

| Metric | MSE (↓) | | | | PCC (↑) | | | |
| --- | --- | --- | --- | --- | --- | --- | --- | --- |
|  | LC2024 (ARPE19) | LC2024 (B16) | LC2024 (N2a) | LC2024 (PC3) | LC2024 (ARPE19) | LC2024 (B16) | LC2024 (N2a) | LC2024 (PC3) |
| AGILE-Metadeta | 2.4359 ± 0.1597 | 2.6538 ± 0.3008 | 2.7428 ± 0.3376 | 3.3561 ± 0.3849 | 0.5364 ± 0.0594 | 0.5816 ± 0.0348 | 0.4872 ± 0.0653 | 0.5913 ± 0.0319 |
| AGILE-MLP | 2.1809 ± 0.1685 | 2.3719 ± 0.3126 | 2.6853 ± 0.4117 | 2.8947 ± 0.3418 | 0.5718 ± 0.0620 | 0.6275 ± 0.0282 | 0.5384 ± 0.0625 | 0.6227 ± 0.0353 |
| AGILE-COMET | 2.7186 ± 0.1230 | 3.1666 ± 0.3438 | 2.9284 ± 0.3890 | 4.1724 ± 0.3720 | 0.3905 ± 0.0492 | 0.3690 ± 0.0488 | 0.4603 ± 0.0880 | 0.2950 ± 0.0579 |
| STRATA (ours) | 1.5560 ± 0.1752 | 1.2414 ± 0.1789 | 1.4281 ± 0.2137 | 1.8796 ± 0.2653 | 0.7232 ± 0.0442 | 0.8193 ± 0.0206 | 0.7866 ± 0.0389 | 0.7725 ± 0.0317 |

#### Additional Experiments

We provide more experiment results in the Appendix C which includes ablation studies (Appendix C.1), hyperparameter sensitivity analysis C.2, cross-dataset experiment C.5.

## 5 Conclusion

In this work, we proposed **STRATA**, a model that effectively predicts LNP transfection efficiency by implicitly modeling interactions. By aligning a ratio-centric view with a molecule-centric view, STRATA jointly considers the two key factors of formulation-level interaction—molecular structure and ratios—and in this process we introduced *Ratio-induced Positional Embedding* (RiPE), a positional encoding scheme that reflects the nature of ratio variables. Extensive experiments demonstrate that this approach is effective under both unseen molecule and unseen ratio settings, and that the model produces predictions that are sensitive to ratio variations.

### Limitations and Future Work

Our work has several limitations that suggest promising directions for future research. First, STRATA relies solely on formulation-level information, without considering manufacturing process parameters (e.g., mixing speed, flow rate) that also influence LNP properties. Second, transfection efficiency is only one of several objectives in LNP design; extending our framework to simultaneously optimize multiple objectives such as immunogenicity and cytotoxicity would better reflect real-world LNP development. Finally, exploring pretraining strategies for formulation screening could enhance its robustness to unseen ratios and novel molecules.

## Acknowledgments and Disclosure of Funding

This work was supported by the Institute of Information & Communications Technology Planning & Evaluation(IITP) grant funded by the Korea government(MSIT) (RS-2025-02304967, AI Star Fellowship(KAIST)). This work was supported by the National Research Foundation of Korea(NRF) grant funded by the Korea government(MSIT) (RS-2024-00335098). This work was supported by National Research Foundation of Korea(NRF) funded by Ministry of Science and ICT (RS-2022-NR068758).

## Appendix

### A Proofs and Properties of RiPE

We adopt the same notation as in the(main text. In particula)r, *R*(*α*) ∈ SO(2) denotes the standard 2×2 rotation matrix, and *R*^*α*^ = diag 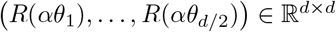 is the block-diagonal rotation operator from Eq. (3) of the main text, parameterised by the frequency schedule Θ = (*θ*_1_, …, *θ*_*d/*2_). Here the superscript *α* is an *index* (the rotation parameter), not a matrix power. The fundamental group property

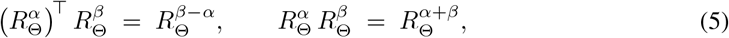

will be used repeatedly.

RiPE encodes a token *x*_*p*_ at position *p* ∈ R_*>*0_ as

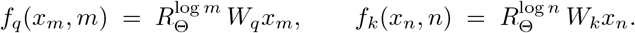

For comparison, RoPE corresponds to the choice *ϕ*_RoPE_(*p*) = *p* and yields ⟨*f*_*q*_(*x*_*m*_, *m*), *f*_*k*_(*x*_*n*_, *n*) ⟩ = *g*(*x*_*m*_, *x*_*n*_, *m* − *n*) [21]; we now show that the RiPE construction transforms this additive structure into a multiplicative one.

#### A.1 Property (i): Ratio Dependence

##### Proposition 2

(Ratio dependence of RiPE). *For all m, n* ∈ R_*>*0_ *and x*_*m*_, *x*_*n*_ ∈ R^*d*^, *there exists a function g such that*

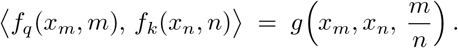

*Proof*. Direct computation using (5) gives

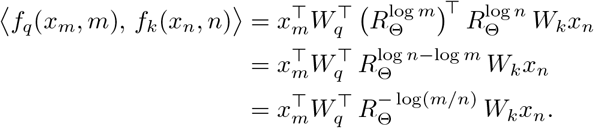

Setting 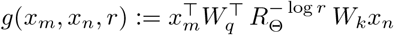 yields the claim.

**A.2 Property (ii): Scale Invariance of Attention Scores**

##### Corollary 1

(Scale invariance). *For all c >* 0, *all m, n* ∈ R_*>*0_, *and all x*_*m*_, *x*_*n*_ ∈ R^*d*^,

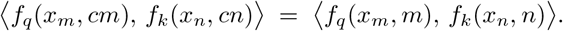

*That is, the attention score is invariant under the diagonal multiplicative rescaling* (*m, n*) ↦ (*cm, cn*).

*Proof*. By Proposition 2, the two sides equal *g x*_*m*_, *x*_*n*_, (*cm*)*/*(*cn*) and *g x*_*m*_, *x*_*n*_, *m/n* respectively. Since (*cm*)*/*(*cn*) = *m/n*, they coincide.

##### Remark 1

(Representation-level equivariance and score-level invariance). *At the level of individual representations, the RiPE encoding transforms equivariantly under the multiplicative group action S*_*c*_ : *p* ↦ *cp: the maps S*_*c*_ *form a group under composition* 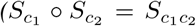, *isomorphic to* (R_*>*0_, ×)*), and a rescaling m* ↦ *cm rotates the encoding* 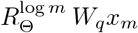 *by an additional factor* 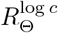. *This per-token rotation, however, cancels in the inner product, yielding the score-level invariance stated in Corollary 1. This is the multiplicative analogue of RoPE’s behaviour under additive translation T*_*c*_ : *p* 1↦ *p* + *c*.

#### A.3 Property (iii): Compositionality of Ratios

We finally show that the rotations associated with pairwise positions are consistent under chaining: traversing *a* → *b* → *c* produces the same rotation as *a* → *c* directly. Define the *relative rotation* between positions *p, q* ∈ R_*>*0_ as

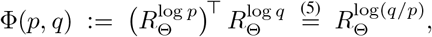

so that 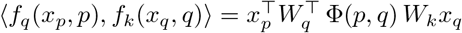.

##### Proposition 3

(Chain composition of ratios). *For all a, b, c* ∈ R_*>*0_,

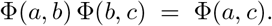

*Equivalently, the ratio along the chain a* → *b* → *c satisfies*

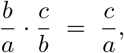

*which is reflected at the level of rotations*.

*Proof*. Using (5),

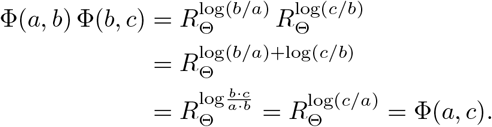

##### Remark 2

(Multiplicativity of ratios). *Proposition 3 is a special case of a more general multiplicative property. Since*

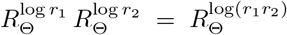

*for any r*_1_, *r*_2_ ∈ R_*>*0_, *the relative rotations of RiPE compose multiplicatively in the ratio: products of pairwise ratios correspond to products of the corresponding rotations. In particular*, Φ(*p, q*) *depends on* (*p, q*) *only through the ratio q/p, so for any a*^*′*^, *b*^*′*^, *c*^*′*^, *d*^*′*^ ∈ R_*>*0_ *we have*

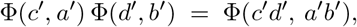

*which mirrors the algebraic identity* 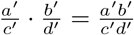 *at the level of representations. Proposition 3 is then obtained as the special case* (*a*^*′*^, *b*^*′*^, *c*^*′*^, *d*^*′*^) = (*b, c, a, b*), *which yields*

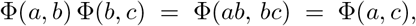

*where the last equality uses bc/*(*ab*) = *c/a*.

##### Corollary 2

(Closure under reversal and identity). *For all a, b* ∈ R_*>*0_,

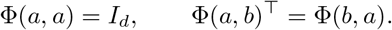

*Proof*. Setting *b* = *a* in 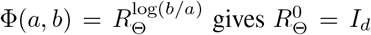. The second identity follows from

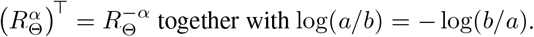

In practical terms, Proposition 3 together with Corollary 2 ensures that RiPE assigns a globally consistent rotation to every pair of positions, so that pairwise ratios extracted by attention can be *transitively* chained without inconsistency—a property naturally aligned with ratio-based reasoning.

#### A.4 Monotonicity and Order Preservation

The previous results characterise the algebraic structure of RiPE; we now record a complementary order-theoretic property, which states that the rotation induced by RiPE reflects the natural ordering of positive ratios.

##### Corollary 3

(Log-linearity and order preservation). *Let r*_1_, *r*_2_ ∈ R_*>*0_. *Then*

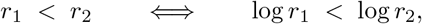

*and consequently, in every frequency block i* = 1, …, *d/*2 *with θ*_*i*_ *>* 0, *the rotation angle is strictly increasing in the ratio:*

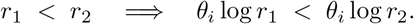

*In particular, the comparison m*_1_*/n* ≶ *m*_2_*/n* ⇔ *m*_1_ ≶ *m*_2_ *is preserved by the relative rotation* Φ(·, ·), *so that distinct ratios induce distinct rotations modulo the intrinsic* 2*π-periodicity of* SO(2).

*Proof*. Strict monotonicity of log on R_*>*0_ gives the first equivalence. The remaining claims follow by multiplying by the (signed) frequency *θ*_*i*_ and applying the definition of Φ.

We note that the 2*π*-periodicity of SO(2) implies that distinct ratios separated by sufficiently large multiplicative factors may be mapped to identical rotations in a given frequency block; the use of multiple frequencies {*θ*_*i*_} mitigates, but does not eliminate, this ambiguity. The order-preservation claim therefore holds locally within each frequency block, and globally only modulo this periodicity. This is the order-theoretic counterpart of the algebraic compositionality established in Proposition 3: ratios closer to 1 produce rotations closer to the identity, and the rotation angle grows (or decays) log-linearly with the ratio.

#### A.5 Summary of Properties

Table 4 consolidates the algebraic and order-theoretic properties of ratios that are reflected by RiPE, together with pointers to the corresponding formal statements.

**Table 4:** Algebraic and order-theoretic properties of ratios, and their realisation in RiPE. Each property is stated symbolically and referenced to its formal counterpart in this appendix.

| # | Property | Statement | Reference |
| --- | --- | --- | --- |
| 1 | Ratio dependence | $\text{score} = g(m/n)$ | Prop. 2 |
| 2 | Scale invariance | $g(cm/cn) = g(m/n)$ | Cor. 1 |
| 3 | Transitivity (chain) | $\frac{b}{a} \cdot \frac{c}{b} = \frac{c}{a}$ | Prop. 3 |
| 4 | Reflexivity (identity) | $\frac{a}{a} = 1, \Phi(a, a) = I$ | Cor. 2 |
| 5 | Antisymmetry / inverse | $\frac{a}{b} = \left(\frac{b}{a}\right)^{-1}$ | Cor. 2 |
| 6 | Multiplicativity | $\frac{ab}{cd} = \frac{a}{c} \cdot \frac{b}{d}$ | Rem. 2 |
| 7 | Log-linearity / monotonicity | $\log(m/n)$ strictly increasing in $m/n$ | Cor. 3 |
| 8 | Order preservation | $m_1/n < m_2/n \iff m_1 < m_2$ | Cor. 3 |

These properties together capture the multiplicative structure of the group (R_*>*0_, ×) (Properties 1–6) and its compatible order (Properties 7–8). This is the precise sense in which RiPE injects a multiplicative inductive bias into the attention geometry: the elementary properties one expects of ratios are reflected in the encoding through the simple device of replacing absolute positions with their logarithms, modulo the 2*π*-periodicity inherent to SO(2).

#### A.6 Further discussions for RiPE

We first note that RiPE is designed to **encode pairwise relationships** between formulation components. At the pairwise level, two components mixed at a 1:2 ratio have the same relative proportion regardless of whether their absolute amounts are 10%:20% or 20%:40%, making an identical pairwise encoding reasonable.

However, one may still question whether the interaction effect should depend on the absolute magnitude of the two components within the full formulation. To make this concern concrete, consider a formulation with *r*_*IL*_ : *r*_*HL*_ : *r*_*CHOL*_ : *r*_*P EG*_ = 10% : 20% : 40% : 30%. In this case, the *r*_*IL*_ : *r*_*HL*_ pair and the *r*_*HL*_ : *r*_*CHOL*_ pair both have a 1:2 ratio and therefore receive the same RiPE encoding. Nevertheless, their overall contributions to the formulation differ substantially: *r*_*IL*_ + *r*_*HL*_ accounts for 30% of the formulation, whereas *r*_*HL*_ + *r*_*CHOL*_ accounts for 60%. Thus, even if their pairwise ratios are identical, one may ask whether treating them identically is appropriate given their different magnitudes in the overall composition.

Importantly, RiPE only defines the **pairwise relational encoding**; it does not by itself determine the final interaction strength. Once the full formulation is processed by the attention mechanism, each interaction is evaluated in the context of all other components. Specifically, for each query, the attention scores are normalized over all keys through softmax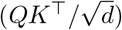. As a result, two pairs with the same RiPE encoding can still receive different final attention weights because they are embedded in different compositional contexts.

Therefore, while RiPE treats identical relative ratios equivalently at the **pairwise** level, their effects can be differentiated at the **formulation** level. In this way, the model can preserve the intended ratio invariance of RiPE while still reflecting differences associated with how strongly each pair contributes to the overall composition.

### B Dataset and Experiment Details

#### B.1 Dataset Details

##### Dataset Source

We surveyed a number of publicly available LNP formulation datasets in order to identify those suitable for our study. Our selection criteria were as follows:

- The dataset must contain multiple candidate molecules for at least one of the four LNP components (IL, HL, CHOL, PEG), and the component ratios must exhibit sufficient diversity across samples.
- The transfection efficiency labels must show a reasonable degree of diversity. Even when the labels are scalar-valued, datasets in which only a handful of distinct values appear across all samples are unsuitable for our purposes.

Based on these criteria, we curated three datasets from the following sources:

- From https://github.com/jswitten/LNP_ML [25], we retained only entries whose Experiment_ID is A549_form_screen, yielding the **A549** dataset. Entries with other experiment IDs did not satisfy the criteria above.
- From https://github.com/evancollins1/LNPDB [8], we retained only entries whose Experiment_ID is YZ_2022, yielding the **YZ2022** dataset. Entries with other experiment IDs did not satisfy the criteria above.
- From https://github.com/MaoResearchGroup/ml_lnp_design_rules [6], the source provides per-cell-type labels for identical LNP formulations. Some cell types have only sparse labeling and therefore yield a small number of samples. We denote the resulting datasets as **LC2024**, where the parenthesized suffix indicates the cell type. Although several cell types satisfy our criteria, including multiple datasets that share identical inputs would be redundant; we therefore selected only two cell types, HEK293 and N2a, for our main experiments, while ARPE19, B16, and PC3 are reserved exclusively for the cross-cell-type generalization experiment.

##### Dataset Characteristics

Each dataset consists of the SMILES strings for the four LNP molecular components (IL, HL, CHOL, PEG), together with the ratios of IL, HL, CHOL, and PEG and an IL-to-cargo ratio. Because the IL becomes positively charged upon ionization and therefore strongly interacts with the oppositely charged nucleic-acid cargo, experimental protocols typically specify the cargo amount relative to the IL rather than as an absolute molar ratio with respect to the cargo. Consequently, while the ratios among the four lipid components are recorded as molar ratios, the IL-to-cargo ratio is reported in alternative units such as the N/P ratio rather than as a molar ratio. As a result, the cargo ratio derived by combining the IL molar ratio with the IL-to-cargo ratio is not a strict molar ratio. Nevertheless, since the true molar ratio cannot be recovered from the available data, and since the reported quantity is itself a meaningful relative ratio defined with respect to a property considered important in LNP formulation, we adopt this derived cargo ratio in our experiments.

The characteristics specific to each dataset are summarized as follows.

- **A549**: contains 192 distinct ILs, 3 HLs, 1 CHOL, and 1 PEG. The dataset was constructed by fixing a small number of non-IL formulations and varying the IL molecule across a large library of candidates. As a result, the diversity of the non-IL components is comparatively limited relative to the other datasets. The IL-to-cargo ratio takes 4 distinct values, although their occurrence frequencies are unevenly distributed. The dataset comprises 1801 samples in total.
- **YZ2022** and **LC2024 (HEK293, B16, PC3)**: each contains 1 IL, 6 HLs, 1 CHOL, and 1 PEG. While molecular diversity is limited, the ratio diversity is rich. Specifically, the dataset was constructed by predefining a set of (IL+HL):(CHOL+PEG) ratios, together with sets of IL:HL and CHOL:PEG ratios, and then enumerating all possible IL:HL:CHOL:PEG combinations from these factors. The IL-to-cargo ratio takes 3 distinct values, each appearing with equal frequency. Each dataset comprises 1080 samples in total.
- **LC2024 (N2a, ARPE19)**: contains 1 IL, 4 HLs, 1 CHOL, and 1 PEG. As with the previous two datasets, molecular diversity is limited but ratio diversity is rich, and the dataset was constructed by enumerating all IL:HL:CHOL:PEG combinations derived from predefined (IL+HL):(CHOL+PEG), IL:HL, and CHOL:PEG ratios. The IL-to-cargo ratio takes 3 distinct values, each appearing with equal frequency. The dataset comprises 720 samples in total.

##### Dataset Split Settings

We construct four split settings: random split, unseen-molecule split, unseen lipid (IL, HL, CHOL, PEG) ratio split, and IL-to-cargo ratio split. Since formulations are typically specified by an IL-to-cargo ratio rather than a cargo molar ratio, we split based on the IL-to-cargo ratio rather than on the cargo ratio, so as to reflect the realistic experimental setting.

We aimed to keep the train/valid/test sample counts consistent across all split types. However, the IL-to-cargo ratio has only a few distinct values, and these values often appear with identical frequencies in the dataset; we therefore use a 1:1:1 ratio for the IL-to-cargo ratio split and a 3:1:1 ratio for all other splits. The characteristics of each split type are summarized below.

- **Unseen-molecule split**: molecules are first grouped by type, and the groups are then partitioned so that no molecule is shared across train, valid, and test sets.
  - **A549**: since IL molecular diversity is high, we split based on IL molecules. The large number of IL types allows a 60%/20%/20% train/valid/test split.
  - **YZ2022** and **LC2024 (HEK293)**: we split based on HL molecules. With 6 HL types available, we assign 1 type to valid, 1 to test, and the remaining 4 to train; the train portion is then downsampled to obtain a 50%/16.7%/16.7% train/valid/test split.
  - **LC2024 (N2a)**: we split based on HL molecules. With 4 HL types available, we assign 1 type to valid, 1 to test, and the remaining 2 to train. In this case, the valid and test portions are downsampled to obtain a 50%/16.7%/16.7% train/valid/test split.
- **Random split**: samples are partitioned uniformly at random without any structural criterion. The train/valid/test proportions follow those of the unseen-molecule split. For LC2024 dataset, cell type ARPE19 follows the setting for N2a and cell type B16, PC3 follows the setting for HEK293.
- **Unseen lipid-component split**: samples are first sorted in ascending order of a chosen lipid component ratio. A contiguous block of size equal to the desired test size is taken as the test set; the remaining samples are then re-sorted, and a contiguous block of size equal to the desired validation size is taken as the validation set. This ensures that the ratio ranges of the three subsets do not overlap. The train/valid/test proportions match those of the unseen-molecule split.
- **IL-to-cargo ratio split**: as noted above, most datasets contain only 3 distinct IL-to-cargo ratios with identical occurrence frequencies, which forces a 1:1:1 split.
  - **A549**: aggregating all samples that share the least frequent IL-to-cargo ratio yields 403 samples. We accordingly assign 400 samples each to train, valid, and test, corresponding to a 22.2%/22.2%/22.2% split of the full dataset.
  - **YZ2022, LC2024 (N2a)**, and **LC2024 (HEK293)**: we assign a different IL-to-cargo ratio value to each of train, valid, and test, yielding a 33.3%/33.3%/33.3% split.

#### B.2 Experiment Setting Details

##### Implementation Details

For the experiments in Appendix C.6, the baseline models LiON, LIFT, and COMET were used exactly as provided by their original implementations. COMET originally comprises two models. The first employs a binary ranking objective as the training loss; following the protocol in the original paper, we evaluated performance at the checkpoint that maximizes the Pearson correlation coefficient (PCC), and we denote this model as **COMET (PCC)**. The second uses the standard MSE loss; in this case, we saved the checkpoint at which the loss is minimized, and we denote it as **COMET (MSE)**.

We describe below the implementation details of the models used in our main experiments, all of which adopt AGILE as the molecule encoder.

- **AGILE-Metadata**: This model follows the architecture illustrated in Figure 2a and is designed with reference to LiON [25]. A learnable AGILE encoder is applied only to the IL to obtain its molecule embedding, while HL, CHOL, and PEG are represented via one-hot encodings. The five ratio values are collected into a 5-dimensional vector. The IL molecule embedding, the three one-hot encodings, and the ratio vector are concatenated and passed through a 2-layer MLP to produce the final prediction.
- **AGILE-MLP**: This model can be viewed as a variant of AGILE-Metadata in which all four components are encoded via AGILE rather than only the IL. Specifically, learnable AGILE encoders are applied to all four components to obtain their molecule embeddings, which are then concatenated with the 5-dimensional ratio vector and passed through a 2-layer MLP.
- **AGILE-COMET**: At a high level, this model replaces the molecule encoder of COMET (MSE) with AGILE. The molecule embeddings of IL, HL, CHOL, and PEG are obtained from a frozen AGILE encoder, and the corresponding lipid ratios are embedded using Gaussian basis functions. Each molecule embedding is concatenated with its associated ratio embedding and projected via a fully connected layer to form a molecule token. The IL-to-cargo ratio is similarly embedded with a Gaussian basis function; however, since no cargo-side information is used (as in the original COMET paper), this ratio embedding alone is projected via a fully connected layer to form a separate token. Together with a learnable CLS token, a total of 6 tokens are fed into a transformer. Only the transformer output corresponding to the CLS token is passed through a 2-layer MLP to produce the prediction; the remaining outputs are discarded.

##### Hyperparameter Search Space

The original baseline models (COMET, LiON, and LIFT) typically report only their best-performing hyperparameters and do not specify a predefined search space. Accordingly, we constructed our search space around each baseline’s reported optimal configuration by adding both smaller and larger candidate values. The search was focused primarily on dimension sizes, the number of attention heads in the multi-head attention layers, and the learning rate. All other hyperparameters not mentioned here follow the original implementation of each model.

The hyperparameter search spaces for the baseline and STRATA are summarized below.

- **STRATA** : The learning rate was searched over {1e-4, 1e-5, 1e-6}, and the hidden dimension over {512, 768, 1024}, with the per-head dimension fixed at 64 (e.g., a hidden dimension of 512 corresponds to 8 attention heads). The alignment loss weight *α* was searched over {0.5, 1.0, 1.5}. We used a weight decay of 0.01, a dropout rate of 0.3, and 4 layers for the Interaction Transformer. For batch training, STRATA was trained with a batch size of 50 and trained for 500 epochs. A cosine annealing learning rate scheduler was not employed, as the model is a relatively small transformer. Since we adopt the pretrained molecule encoder from AGILE, its hyperparameters follow the original configuration: a node embedding dimension of 300 and 5 layers.
- **AGILE-Metadata**: Hyperparameters related to the molecule encoder follow AGILE [28], while the remaining hyperparameters follow LiON [25]. The searched values are: learning rate ∈ {10^*−*4^, 10^*−*5^, 10^*−*6^} and hidden dimension ∈ {128, 256, 512}.
- **AGILE-MLP**: All hyperparameters generally follow AGILE [28]. The searched values are: learning rate ∈ {10^*−*4^, 10^*−*5^, 10^*−*6^} and hidden dimension ∈ {128, 256, 512}.
- **AGILE-COMET**: Hyperparameters related to the molecule encoder follow AGILE [28], while the remaining hyperparameters follow COMET [4]. Since the original COMET paper reports 8 transformer layers as the best configuration, we searched neighboring values. The searched values are: number of transformer layers ∈ {6, 8, 10, transformer hidden dimension ∈ {256, 512}, number of attention heads ∈ {4, 8, 16}, and learning rate ∈ {10^*−*3^, 10^*−*4^, 10^*−*5^}. Note that the Gaussian-basis-function ratio embedding uses 128 dimensions, identical to the original COMET implementation.
- **COMET**: Hyperparameters generally follow the best configuration reported in the original paper. The searched values are: number of transformer layers ∈ {6, 8, 10} (centered on the reported best of 8), transformer hidden dimension ∈ {256, 512}, number of attention heads ∈ {4, 8, 16}, and learning rate ∈ {10^*−*3^, 10^*−*4^, 10^*−*5^}. As above, the Gaussian-basis-function ratio embedding uses 128 dimensions, identical to the original COMET implementation.
- **LIFT**: Hyperparameters generally follow the best configuration reported in the original paper. The searched values are: learning rate ∈ {10^*−*4^, 10^*−*5^, 10^*−*6^}, hidden dimension ∈ {128, 256, 512}, number of attention heads ∈ {4, 8, 16}, and number of GNN layers ∈ {1, 2, 3}.
- **LiON**: Hyperparameters generally follow the best configuration reported in the original paper. The searched values are: learning rate ∈ {10^*−*4^, 10^*−*5^, 10^*−*6^}, hidden dimension ∈ {128, 256, 512}, and number of GNN layers ∈ {3, 4, 5}. Note that the original LiON implementation normalizes the metadata features. Although that work normalized the metadata due to the large number of metadata fields, in our setting the number of metadata fields is small and the scale gap between the one-hot encodings and the ratio values is large; we therefore disable the normalization.

#### B.3 Experiments compute resource

We used a single 48GB NVIDIA RTX A6000 GPU in Ubuntu environment.

### C Additional Experiments

#### C.1 Ablation Studies

We conduct a comprehensive ablation study to assess the contribution of the alignment loss *L*_align_ to model performance. Each row pair in Table 5 contrasts the variant trained *without L*_align_ (✗) against the full model trained *with L*_align_ (❍), evaluated on four datasets (A549, YZ2022, LC2024 (N2a), LC2024 (HEK293)) under seven different split setting identical to the main experiment. We report mean and standard deviation over ten random seeds.

**Table 5:** Ablation study for investigating the impact of alignment loss.

| Ablation<br>$\mathcal{L}_{\text{align}}$ | MSE (↓) | | | | PCC (↑) | | | |
| --- | --- | --- | --- | --- | --- | --- | --- | --- |
|  | A549 | YZ2022 | LC2024 (N2a) | LC2024 (HEK293) | A549 | YZ2022 | LC2024 (N2a) | LC2024 (HEK293) |
|  | random split |  |  |  |  |  |  |  |
| X | 0.5217 ± 0.0297 | 0.3897 ± 0.0682 | 0.8344 ± 0.1589 | 2.7368 ± 0.4227 | 0.7903 ± 0.0152 | 0.7870 ± 0.0657 | 0.8847 ± 0.0254 | 0.7066 ± 0.0509 |
| O | 0.4873 ± 0.0290 | 0.3698 ± 0.0667 | 0.7953 ± 0.1295 | 2.5155 ± 0.4417 | 0.8030 ± 0.0128 | 0.8015 ± 0.0452 | 0.8863 ± 0.0225 | 0.7269 ± 0.0558 |
|  | unseen molecule split |  |  |  |  |  |  |  |
| X | 0.6973 ± 0.1291 | 1.0850 ± 0.5998 | 3.5464 ± 2.0736 | 5.4797 ± 2.0478 | 0.7216 ± 0.0396 | 0.1751 ± 0.2645 | 0.4818 ± 0.2531 | 0.4420 ± 0.1394 |
| O | 0.6558 ± 0.0966 | 0.8652 ± 0.3153 | 2.8728 ± 1.1652 | 4.5784 ± 0.8397 | 0.7415 ± 0.0326 | 0.3451 ± 0.2211 | 0.5051 ± 0.1682 | 0.5209 ± 0.0977 |
|  | unseen IL ratio split |  |  |  |  |  |  |  |
| X | 0.6507 ± 0.0780 | 0.7012 ± 0.3642 | 1.6567 ± 0.5821 | 4.1501 ± 1.0806 | 0.7012 ± 0.0675 | 0.6479 ± 0.1271 | 0.7621 ± 0.1020 | 0.4679 ± 0.1415 |
| O | 0.6391 ± 0.0909 | 0.5487 ± 0.2989 | 1.5142 ± 0.5878 | 3.5891 ± 0.8104 | 0.7236 ± 0.0837 | 0.7142 ± 0.1136 | 0.7636 ± 0.1317 | 0.5392 ± 0.1484 |
|  | unseen HL ratio split |  |  |  |  |  |  |  |
| X | 0.7383 ± 0.1014 | 0.6840 ± 0.4165 | 1.7694 ± 1.9619 | 3.3519 ± 0.9994 | 0.7171 ± 0.0477 | 0.4963 ± 0.0822 | 0.5908 ± 0.1871 | 0.5860 ± 0.0882 |
| O | 0.6807 ± 0.0461 | 0.4757 ± 0.1957 | 1.6280 ± 1.9178 | 3.2386 ± 0.8974 | 0.7288 ± 0.0332 | 0.5404 ± 0.0801 | 0.5963 ± 0.1698 | 0.5983 ± 0.0906 |
|  | unseen CHOL ratio split |  |  |  |  |  |  |  |
| X | 0.7438 ± 0.0915 | 0.4833 ± 0.1654 | 1.3521 ± 0.4647 | 4.2758 ± 1.6356 | 0.6802 ± 0.0468 | 0.7671 ± 0.0535 | 0.8242 ± 0.0630 | 0.5113 ± 0.1814 |
| O | 0.6290 ± 0.0987 | 0.4084 ± 0.0934 | 1.3402 ± 0.4061 | 3.4368 ± 1.2848 | 0.7245 ± 0.0435 | 0.7965 ± 0.0498 | 0.8258 ± 0.0557 | 0.5823 ± 0.1571 |
|  | unseen PEG ratio split |  |  |  |  |  |  |  |
| X | 0.7886 ± 0.1474 | 0.5910 ± 0.1167 | 1.2843 ± 0.4417 | 4.3332 ± 1.5343 | 0.6297 ± 0.0544 | 0.7078 ± 0.0921 | 0.7788 ± 0.0835 | 0.3732 ± 0.1900 |
| O | 0.6694 ± 0.1668 | 0.5660 ± 0.0937 | 1.1286 ± 0.1761 | 3.9178 ± 0.9480 | 0.7121 ± 0.0481 | 0.7441 ± 0.0767 | 0.8069 ± 0.0303 | 0.3551 ± 0.2069 |
|  | unseen IL-to-cargo ratio split |  |  |  |  |  |  |  |
| X | 1.4596 ± 0.2728 | 0.5172 ± 0.1152 | 1.4020 ± 0.3429 | 4.1384 ± 1.4423 | 0.4558 ± 0.0828 | 0.7373 ± 0.0507 | 0.7873 ± 0.0780 | 0.5935 ± 0.1180 |
| O | 1.3575 ± 0.2834 | 0.4735 ± 0.1012 | 1.3321 ± 0.2225 | 3.9980 ± 1.3497 | 0.4910 ± 0.1011 | 0.7524 ± 0.0395 | 0.8084 ± 0.0402 | 0.6120 ± 0.1040 |

Across all 56 dataset–metric–split combinations, incorporating *L*_align_ improves performance in 55 cases, with the sole exception being PCC on LC2024 (HEK293) under the unseen PEG-ratio split, where the two variants are statistically indistinguishable (0.3732 ± 0.1900 vs. 0.3551 ± 0.2069). As shown in Table 1, however, not only STRATA but all baselines exhibit degraded performance on this particular split, suggesting that this setting is intrinsically difficult to fit regardless of whether the alignment loss is applied. This consistent improvement across heterogeneous datasets and split protocols indicates that the gain from *L*_align_ is robust rather than dataset-specific. Taken together, these ablations support our central claim that *L*_align_ is not merely a regularizer for in-distribution performance but a key driver of out-of-distribution generalization across unseen molecules and unseen compositional ratios.

To validate the necessity of the molecule-centric view, we conducted an ablation study using a model that relies solely on the ratio-centric view. In this variant, predictions were made directly from the output of the ratio-centric view. Since the molecule-centric view was removed, the alignment loss (*L*_align_) was also excluded.

Table 6 presents the results. As shown in the table, this variant performs worse than the full STRATA model. Although molecular interactions can alter the representation of each component, their effects should remain grounded in the underlying molecular structure and identity. Removing the molecule-centric route weakens this molecule-specific grounding, demonstrating that RiPE alone is insufficient to account for the performance gains achieved by the full architecture.

**Table 6:** Ablation study for investigating the impact of molecule-centric view.

| Ablation<br>molecule-centric view | MSE (↓) | | | | PCC (↑) | | | |
| --- | --- | --- | --- | --- | --- | --- | --- | --- |
|  | A549 | YZ2022 | LC2024 (N2a) | LC2024 (HEK293) | A549 | YZ2022 | LC2024 (N2a) | LC2024 (HEK293) |
|  | random split |  |  |  |  |  |  |  |
| X | 0.7659 ± 0.0392 | 0.5683 ± 0.0833 | 2.0596 ± 0.5755 | 3.6670 ± 0.4861 | 0.6587 ± 0.0169 | 0.6666 ± 0.0523 | 0.6893 ± 0.0928 | 0.5624 ± 0.0506 |
| O | 0.4873 ± 0.0290 | 0.3698 ± 0.0667 | 0.7953 ± 0.1295 | 2.5155 ± 0.4417 | 0.8030 ± 0.0128 | 0.8015 ± 0.0452 | 0.8863 ± 0.0225 | 0.7269 ± 0.0558 |
|  | unseen molecule split |  |  |  |  |  |  |  |
| X | 0.8124 ± 0.1507 | 1.0916 ± 0.4189 | 3.0475 ± 1.0214 | 5.5247 ± 1.3832 | 0.6537 ± 0.0546 | 0.1579 ± 0.2324 | 0.5337 ± 0.1872 | 0.4580 ± 0.1640 |
| O | 0.6558 ± 0.0966 | 0.8652 ± 0.3153 | 2.8728 ± 1.1652 | 4.5784 ± 0.8397 | 0.7415 ± 0.0326 | 0.3451 ± 0.2211 | 0.5051 ± 0.1682 | 0.5209 ± 0.0977 |
|  | unseen IL ratio split |  |  |  |  |  |  |  |
| X | 0.7969 ± 0.1109 | 0.6145 ± 0.2254 | 3.0719 ± 1.5011 | 4.3186 ± 0.2147 | 0.5837 ± 0.1057 | 0.6210 ± 0.1001 | 0.6022 ± 0.0931 | 0.4276 ± 0.1729 |
| O | 0.6391 ± 0.0909 | 0.5487 ± 0.2989 | 1.5142 ± 0.5878 | 3.5891 ± 0.8104 | 0.7236 ± 0.0837 | 0.7142 ± 0.1136 | 0.7636 ± 0.1317 | 0.5392 ± 0.1484 |
|  | unseen HL ratio split |  |  |  |  |  |  |  |
| X | 1.1100 ± 0.2549 | 0.4985 ± 0.1669 | 2.1285 ± 3.4696 | 3.2762 ± 0.4669 | 0.5242 ± 0.1348 | 0.4780 ± 0.0852 | 0.5941 ± 0.1939 | 0.5382 ± 0.0665 |
| O | 0.6807 ± 0.0461 | 0.4757 ± 0.1957 | 1.6280 ± 1.9178 | 3.2386 ± 0.8974 | 0.7288 ± 0.0332 | 0.5404 ± 0.0801 | 0.5963 ± 0.1698 | 0.5983 ± 0.0906 |
|  | unseen CHOL ratio split |  |  |  |  |  |  |  |
| X | 0.8904 ± 0.1666 | 0.6192 ± 0.1473 | 2.2438 ± 0.6563 | 4.3476 ± 1.7447 | 0.5571 ± 0.0949 | 0.6619 ± 0.0652 | 0.6514 ± 0.0919 | 0.4805 ± 0.1267 |
| O | 0.6290 ± 0.0987 | 0.4084 ± 0.0934 | 1.3402 ± 0.4061 | 3.4368 ± 1.2848 | 0.7245 ± 0.0435 | 0.7965 ± 0.0498 | 0.8258 ± 0.0557 | 0.5823 ± 0.1571 |
|  | unseen PEG ratio split |  |  |  |  |  |  |  |
| X | 1.1083 ± 0.3146 | 0.7122 ± 0.1776 | 2.5007 ± 0.7831 | 4.7829 ± 1.8566 | 0.4447 ± 0.1814 | 0.6161 ± 0.0748 | 0.4728 ± 0.1920 | 0.1373 ± 0.2333 |
| O | 0.6694 ± 0.1668 | 0.5660 ± 0.0937 | 1.1286 ± 0.1761 | 3.9178 ± 0.9480 | 0.7121 ± 0.0481 | 0.7441 ± 0.0767 | 0.8069 ± 0.0303 | 0.3551 ± 0.2069 |
|  | unseen IL-to-cargo ratio split |  |  |  |  |  |  |  |
| X | 1.6554 ± 0.1951 | 0.5720 ± 0.1042 | 1.7704 ± 0.2762 | 4.2340 ± 1.0128 | 0.1405 ± 0.0426 | 0.6832 ± 0.0410 | 0.7520 ± 0.0426 | 0.5497 ± 0.0502 |
| O | 1.3575 ± 0.2834 | 0.4735 ± 0.1012 | 1.3321 ± 0.2225 | 3.9980 ± 1.3497 | 0.4910 ± 0.1011 | 0.7524 ± 0.0395 | 0.8084 ± 0.0402 | 0.6120 ± 0.1040 |

#### C.2 Hyperparameter Sensitivity Analysis

To analyze the influence of the principal hyperparameters on STRATA, we vary (i) the alignment loss weight *α* and (ii) the model capacity, controlled jointly by the hidden dimension and the number of attention heads. For each configuration, we report MSE (↓) and PCC (↑) on all four datasets under the random split and the six unseen-component splits, with mean and standard deviation computed over ten random seeds.

##### Sensitivity to the alignment loss weight *α*

Table 7 reports the performance of STRATA for *α* ∈ {0.5, 1.0, 1.5}, where *α* controls the relative contribution of *L*_align_ to the overall training objective. Note that the variant without *L*_align_ in the ablation study (Appendix C.1) is equivalent to setting *α* = 0, and we therefore interpret the two studies jointly as a sweep over *α* ∈ {0, 0.5, 1.0, 1.5}. Under this combined view, performance tends to improve as *α* grows from zero and then deteriorates once *α* becomes too large. This non-monotonic pattern suggests that aligning the two views is beneficial for learning, but that an overly strong alignment penalty can interfere with each view’s ability to exploit its own inductive bias and capture view-specific informative signals.

**Table 7:**
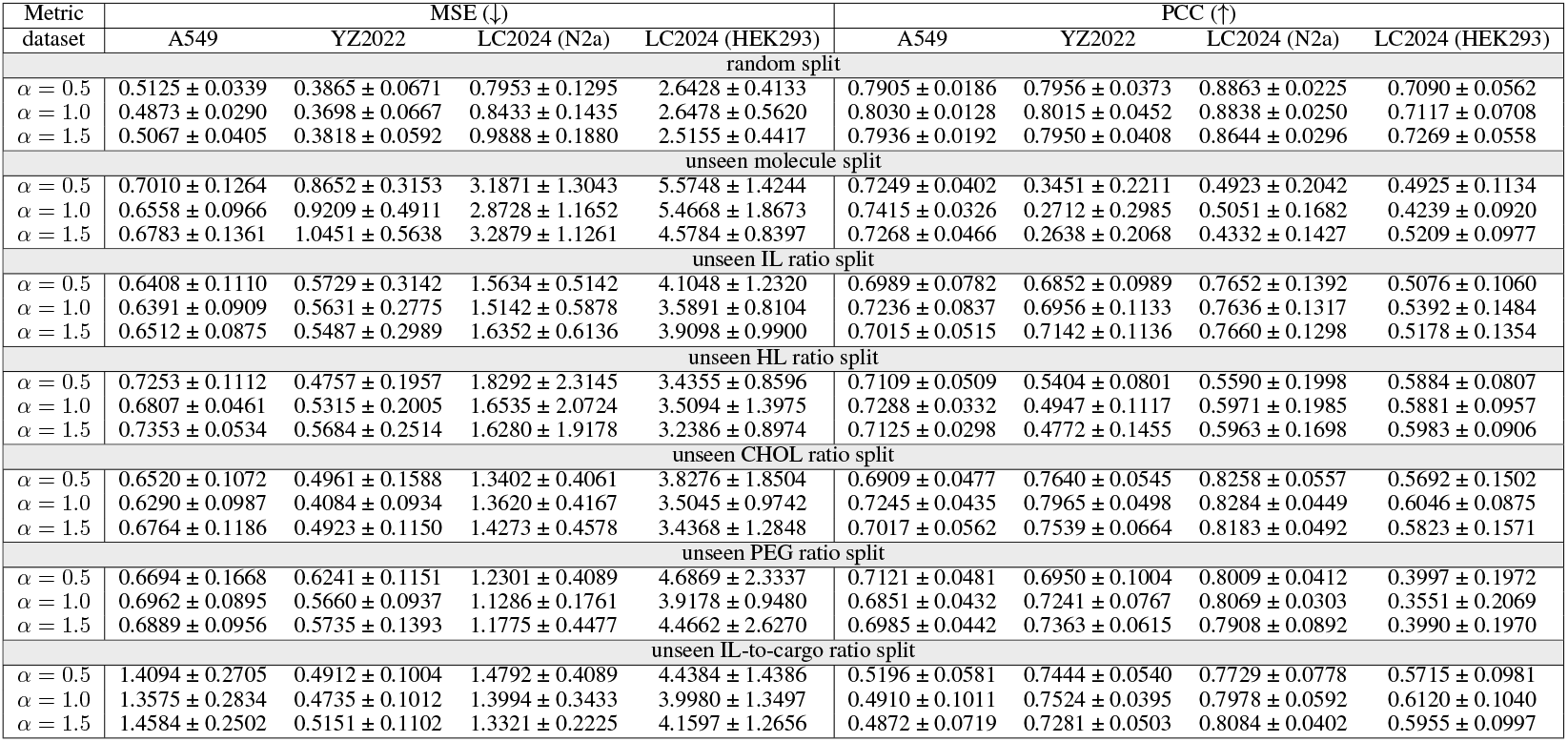
Hyperparameter sensitivity analysis for *α*.

| Metric<br>dataset | MSE (↓) | | | | PCC (↑) | | | |
| --- | --- | --- | --- | --- | --- | --- | --- | --- |
|  | A549 | YZ2022 | LC2024 (N2a) | LC2024 (HEK293) | A549 | YZ2022 | LC2024 (N2a) | LC2024 (HEK293) |
| random split |  |  |  |  |  |  |  |  |
| $\alpha = 0.5$ | 0.5125 ± 0.0339 | 0.3865 ± 0.0671 | 0.7953 ± 0.1295 | 2.6428 ± 0.4133 | 0.7905 ± 0.0186 | 0.7956 ± 0.0373 | 0.8863 ± 0.0225 | 0.7090 ± 0.0562 |
| $\alpha = 1.0$ | 0.4873 ± 0.0290 | 0.3698 ± 0.0667 | 0.8433 ± 0.1435 | 2.6478 ± 0.5620 | 0.8030 ± 0.0128 | 0.8015 ± 0.0452 | 0.8838 ± 0.0250 | 0.7117 ± 0.0708 |
| $\alpha = 1.5$ | 0.5067 ± 0.0405 | 0.3818 ± 0.0592 | 0.9888 ± 0.1880 | 2.5155 ± 0.4417 | 0.7936 ± 0.0192 | 0.7950 ± 0.0408 | 0.8644 ± 0.0296 | 0.7269 ± 0.0558 |
| unseen molecule split |  |  |  |  |  |  |  |  |
| $\alpha = 0.5$ | 0.7010 ± 0.1264 | 0.8652 ± 0.3153 | 3.1871 ± 1.3043 | 5.5748 ± 1.4244 | 0.7249 ± 0.0402 | 0.3451 ± 0.2211 | 0.4923 ± 0.2042 | 0.4925 ± 0.1134 |
| $\alpha = 1.0$ | 0.6558 ± 0.0966 | 0.9209 ± 0.4911 | 2.8728 ± 1.1652 | 5.4668 ± 1.8673 | 0.7415 ± 0.0326 | 0.2712 ± 0.2985 | 0.5051 ± 0.1682 | 0.4239 ± 0.0920 |
| $\alpha = 1.5$ | 0.6783 ± 0.1361 | 1.0451 ± 0.5638 | 3.2879 ± 1.1261 | 4.5784 ± 0.8397 | 0.7268 ± 0.0466 | 0.2638 ± 0.2068 | 0.4332 ± 0.1427 | 0.5209 ± 0.0977 |
| unseen IL ratio split |  |  |  |  |  |  |  |  |
| $\alpha = 0.5$ | 0.6408 ± 0.1110 | 0.5729 ± 0.3142 | 1.5634 ± 0.5142 | 4.1048 ± 1.2320 | 0.6989 ± 0.0782 | 0.6852 ± 0.0989 | 0.7652 ± 0.1392 | 0.5076 ± 0.1060 |
| $\alpha = 1.0$ | 0.6391 ± 0.0909 | 0.5631 ± 0.2775 | 1.5142 ± 0.5878 | 3.5891 ± 0.8104 | 0.7236 ± 0.0837 | 0.6956 ± 0.1133 | 0.7636 ± 0.1317 | 0.5392 ± 0.1484 |
| $\alpha = 1.5$ | 0.6512 ± 0.0875 | 0.5487 ± 0.2989 | 1.6352 ± 0.6136 | 3.9098 ± 0.9900 | 0.7015 ± 0.0515 | 0.7142 ± 0.1136 | 0.7660 ± 0.1298 | 0.5178 ± 0.1354 |
| unseen HL ratio split |  |  |  |  |  |  |  |  |
| $\alpha = 0.5$ | 0.7253 ± 0.1112 | 0.4757 ± 0.1957 | 1.8292 ± 2.3145 | 3.4355 ± 0.8596 | 0.7109 ± 0.0509 | 0.5404 ± 0.0801 | 0.5590 ± 0.1998 | 0.5884 ± 0.0807 |
| $\alpha = 1.0$ | 0.6807 ± 0.0461 | 0.5315 ± 0.2005 | 1.6535 ± 2.0724 | 3.5094 ± 1.3975 | 0.7288 ± 0.0332 | 0.4947 ± 0.1117 | 0.5971 ± 0.1985 | 0.5881 ± 0.0957 |
| $\alpha = 1.5$ | 0.7353 ± 0.0534 | 0.5684 ± 0.2514 | 1.6280 ± 1.9178 | 3.2386 ± 0.8974 | 0.7125 ± 0.0298 | 0.4772 ± 0.1455 | 0.5963 ± 0.1698 | 0.5983 ± 0.0906 |
| unseen CHOL ratio split |  |  |  |  |  |  |  |  |
| $\alpha = 0.5$ | 0.6520 ± 0.1072 | 0.4961 ± 0.1588 | 1.3402 ± 0.4061 | 3.8276 ± 1.8504 | 0.6909 ± 0.0477 | 0.7640 ± 0.0545 | 0.8258 ± 0.0557 | 0.5692 ± 0.1502 |
| $\alpha = 1.0$ | 0.6290 ± 0.0987 | 0.4084 ± 0.0934 | 1.3620 ± 0.4167 | 3.5045 ± 0.9742 | 0.7245 ± 0.0435 | 0.7965 ± 0.0498 | 0.8284 ± 0.0449 | 0.6046 ± 0.0875 |
| $\alpha = 1.5$ | 0.6764 ± 0.1186 | 0.4923 ± 0.1150 | 1.4273 ± 0.4578 | 3.4368 ± 1.2848 | 0.7017 ± 0.0562 | 0.7539 ± 0.0664 | 0.8183 ± 0.0492 | 0.5823 ± 0.1571 |
| unseen PEG ratio split |  |  |  |  |  |  |  |  |
| $\alpha = 0.5$ | 0.6694 ± 0.1668 | 0.6241 ± 0.1151 | 1.2301 ± 0.4089 | 4.6869 ± 2.3337 | 0.7121 ± 0.0481 | 0.6950 ± 0.1004 | 0.8009 ± 0.0412 | 0.3997 ± 0.1972 |
| $\alpha = 1.0$ | 0.6962 ± 0.0895 | 0.5660 ± 0.0937 | 1.1286 ± 0.1761 | 3.9178 ± 0.9480 | 0.6851 ± 0.0432 | 0.7241 ± 0.0767 | 0.8069 ± 0.0303 | 0.3551 ± 0.2069 |
| $\alpha = 1.5$ | 0.6889 ± 0.0956 | 0.5735 ± 0.1393 | 1.1775 ± 0.4477 | 4.4662 ± 2.6270 | 0.6985 ± 0.0442 | 0.7363 ± 0.0615 | 0.7908 ± 0.0892 | 0.3990 ± 0.1970 |
| unseen IL-to-cargo ratio split |  |  |  |  |  |  |  |  |
| $\alpha = 0.5$ | 1.4094 ± 0.2705 | 0.4912 ± 0.1004 | 1.4792 ± 0.4089 | 4.4384 ± 1.4386 | 0.5196 ± 0.0581 | 0.7444 ± 0.0540 | 0.7729 ± 0.0778 | 0.5715 ± 0.0981 |
| $\alpha = 1.0$ | 1.3575 ± 0.2834 | 0.4735 ± 0.1012 | 1.3994 ± 0.3433 | 3.9980 ± 1.3497 | 0.4910 ± 0.1011 | 0.7524 ± 0.0395 | 0.7978 ± 0.0592 | 0.6120 ± 0.1040 |
| $\alpha = 1.5$ | 1.4584 ± 0.2502 | 0.5151 ± 0.1102 | 1.3321 ± 0.2225 | 4.1597 ± 1.2656 | 0.4872 ± 0.0719 | 0.7281 ± 0.0503 | 0.8084 ± 0.0402 | 0.5955 ± 0.0997 |

##### Sensitivity to model capacity

Table 8 examines three architecture configurations of increasing capacity: (512 dim, 8 heads), (768 dim, 12 heads), and (1024 dim, 16 heads). The differences across configurations are typically within one standard deviation of the seed-level variability, and the best-performing setting fluctuates across datasets and splits without a single configuration dominating universally, indicating that model capacity is not a critical factor for STRATA’s performance.

**Table 8:** Hyperparameter sensitivity analysis for hidden dimension paired with number of heads.

| Metric<br>dataset | MSE (↓) | | | | PCC (↑) | | | |
| --- | --- | --- | --- | --- | --- | --- | --- | --- |
|  | A549 | YZ2022 | LC2024 (N2a) | LC2024 (HEK293) | A549 | YZ2022 | LC2024 (N2a) | LC2024 (HEK293) |
| random split |  |  |  |  |  |  |  |  |
| 512 dim, 8 head | 0.4873 ± 0.0290 | 0.3747 ± 0.0592 | 0.8980 ± 0.2445 | 2.7965 ± 0.4429 | 0.8030 ± 0.0128 | 0.7946 ± 0.0318 | 0.8786 ± 0.0330 | 0.6977 ± 0.0645 |
| 768 dim, 12 head | 0.5080 ± 0.0332 | 0.3934 ± 0.0761 | 0.8448 ± 0.1610 | 2.6786 ± 0.6082 | 0.7939 ± 0.0130 | 0.7866 ± 0.0521 | 0.8868 ± 0.0214 | 0.7243 ± 0.0663 |
| 1024 dim, 16 head | 0.5192 ± 0.0346 | 0.3698 ± 0.0667 | 0.7953 ± 0.1295 | 2.5155 ± 0.4417 | 0.7897 ± 0.0146 | 0.8015 ± 0.0452 | 0.8863 ± 0.0225 | 0.7269 ± 0.0558 |
| unseen molecule split |  |  |  |  |  |  |  |  |
| 512 dim, 8 head | 0.6558 ± 0.0966 | 0.9399 ± 0.3790 | 3.5002 ± 1.4257 | 4.5784 ± 0.8397 | 0.7415 ± 0.0326 | 0.2679 ± 0.2041 | 0.4496 ± 0.2515 | 0.5209 ± 0.0977 |
| 768 dim, 12 head | 0.6995 ± 0.0971 | 0.9520 ± 0.4840 | 3.1487 ± 1.3224 | 5.6620 ± 1.7334 | 0.7239 ± 0.0317 | 0.2794 ± 0.2055 | 0.5666 ± 0.0955 | 0.4873 ± 0.1166 |
| 1024 dim, 16 head | 0.7171 ± 0.1157 | 0.8652 ± 0.3153 | 2.8728 ± 1.1652 | 5.6750 ± 2.1701 | 0.7176 ± 0.0375 | 0.3451 ± 0.2211 | 0.5051 ± 0.1682 | 0.4343 ± 0.1403 |
| unseen IL ratio split |  |  |  |  |  |  |  |  |
| 512 dim, 8 head | 0.6391 ± 0.0909 | 0.6835 ± 0.4295 | 1.9541 ± 0.7920 | 4.3223 ± 1.0742 | 0.7236 ± 0.0837 | 0.6618 ± 0.0898 | 0.7135 ± 0.1055 | 0.4654 ± 0.0855 |
| 768 dim, 12 head | 0.6520 ± 0.1166 | 0.5487 ± 0.2989 | 1.5142 ± 0.5878 | 3.7936 ± 0.6827 | 0.6894 ± 0.0881 | 0.7142 ± 0.1136 | 0.7636 ± 0.1317 | 0.5134 ± 0.1183 |
| 1024 dim, 16 head | 0.6472 ± 0.1129 | 0.6082 ± 0.2814 | 1.6615 ± 0.6179 | 3.5891 ± 0.8104 | 0.7031 ± 0.0727 | 0.6729 ± 0.1512 | 0.7389 ± 0.1661 | 0.5392 ± 0.1484 |
| unseen HL ratio split |  |  |  |  |  |  |  |  |
| 512 dim, 8 head | 0.6807 ± 0.0461 | 0.5565 ± 0.2112 | 1.8710 ± 1.9944 | 4.1750 ± 2.0499 | 0.7288 ± 0.0332 | 0.4956 ± 0.0832 | 0.5540 ± 0.1954 | 0.5271 ± 0.1199 |
| 768 dim, 12 head | 0.7551 ± 0.1624 | 0.5199 ± 0.2138 | 1.7025 ± 2.1781 | 4.2015 ± 1.9634 | 0.7186 ± 0.0665 | 0.5264 ± 0.0993 | 0.6081 ± 0.1946 | 0.5562 ± 0.1111 |
| 1024 dim, 16 head | 0.7251 ± 0.1007 | 0.4757 ± 0.1957 | 1.6280 ± 1.9178 | 3.2386 ± 0.8974 | 0.7195 ± 0.0577 | 0.5404 ± 0.0801 | 0.5963 ± 0.1698 | 0.5983 ± 0.0906 |
| unseen CHOL ratio split |  |  |  |  |  |  |  |  |
| 512 dim, 8 head | 0.6290 ± 0.0987 | 0.5244 ± 0.1679 | 1.5325 ± 0.3791 | 4.1040 ± 0.8251 | 0.7245 ± 0.0435 | 0.7427 ± 0.0764 | 0.7636 ± 0.0733 | 0.5137 ± 0.0936 |
| 768 dim, 12 head | 0.6468 ± 0.0946 | 0.4084 ± 0.0934 | 1.3726 ± 0.5351 | 3.8262 ± 0.9833 | 0.7006 ± 0.0469 | 0.7965 ± 0.0498 | 0.8228 ± 0.0616 | 0.5401 ± 0.1842 |
| 1024 dim, 16 head | 0.6553 ± 0.1048 | 0.5778 ± 0.3717 | 1.3402 ± 0.4061 | 3.4368 ± 1.2848 | 0.7026 ± 0.0470 | 0.7389 ± 0.1042 | 0.8258 ± 0.0557 | 0.5823 ± 0.1571 |
| unseen PEG ratio split |  |  |  |  |  |  |  |  |
| 512 dim, 8 head | 0.6694 ± 0.1668 | 0.6002 ± 0.2077 | 1.3506 ± 0.2312 | 4.6649 ± 2.0865 | 0.7121 ± 0.0481 | 0.7086 ± 0.0899 | 0.7488 ± 0.1125 | 0.3679 ± 0.1903 |
| 768 dim, 12 head | 0.7161 ± 0.1222 | 0.5819 ± 0.1753 | 1.1442 ± 0.2627 | 3.9178 ± 0.9480 | 0.6623 ± 0.0567 | 0.7131 ± 0.0884 | 0.8072 ± 0.0418 | 0.3551 ± 0.2069 |
| 1024 dim, 16 head | 0.7365 ± 0.1039 | 0.5660 ± 0.0937 | 1.1286 ± 0.1761 | 4.3240 ± 1.7602 | 0.6553 ± 0.0354 | 0.7241 ± 0.0767 | 0.8069 ± 0.0303 | 0.3710 ± 0.1784 |
| unseen IL-to-cargo ratio split |  |  |  |  |  |  |  |  |
| 512 dim, 8 head | 1.3575 ± 0.2834 | 0.5662 ± 0.1942 | 1.5325 ± 0.3791 | 4.7166 ± 1.6850 | 0.4910 ± 0.1011 | 0.7034 ± 0.1038 | 0.7636 ± 0.0733 | 0.5170 ± 0.1376 |
| 768 dim, 12 head | 1.5733 ± 0.3382 | 0.5071 ± 0.1272 | 1.5408 ± 0.4345 | 4.0768 ± 1.1949 | 0.4526 ± 0.1092 | 0.7392 ± 0.0632 | 0.7797 ± 0.0673 | 0.5790 ± 0.0997 |
| 1024 dim, 16 head | 1.6978 ± 0.5351 | 0.4735 ± 0.1012 | 1.3321 ± 0.2225 | 3.9980 ± 1.3497 | 0.4476 ± 0.0810 | 0.7524 ± 0.0395 | 0.8084 ± 0.0402 | 0.6120 ± 0.1040 |

#### C.3 Time Complexity

To analyze the computational cost of STRATA relative to baselines, we measure the wall-clock time required for a single training epoch ten times per method and report the mean and standard deviation in Table 9.

**Table 9:**
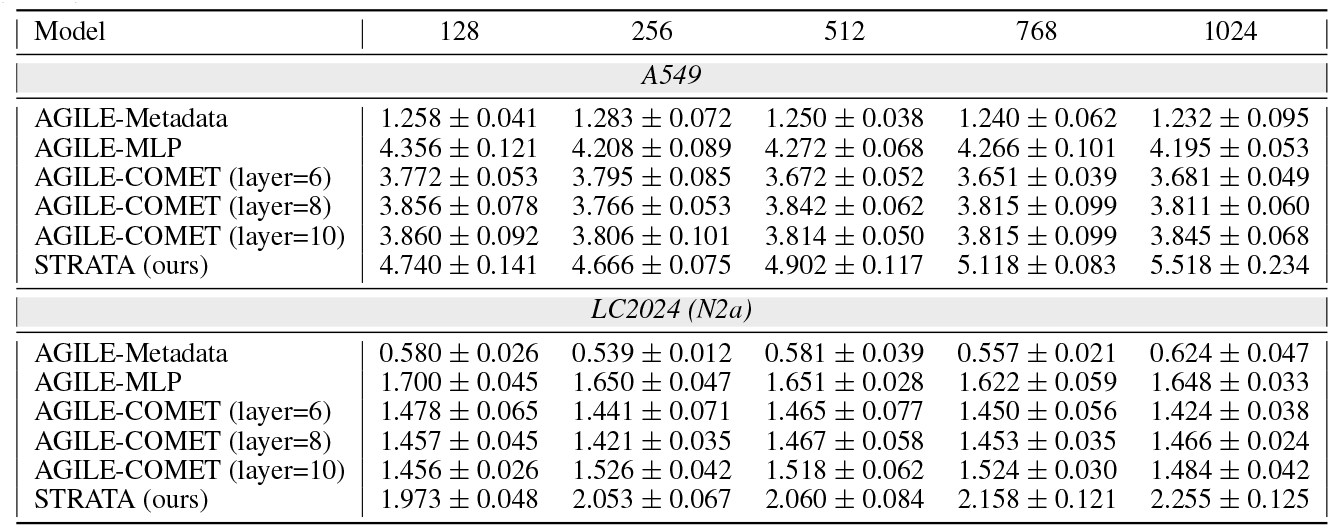
Training time per epoch (in seconds) across different hidden dimensions on A549 and LC2024 (N2a) datasets. The column denotes hidden dimension.

| Model | 128 | 256 | 512 | 768 | 1024 |
| --- | --- | --- | --- | --- | --- |
| *A549* |  |  |  |  |  |
| AGILE-Metaddata | 1.258 ± 0.041 | 1.283 ± 0.072 | 1.250 ± 0.038 | 1.240 ± 0.062 | 1.232 ± 0.095 |
| AGILE-MLP | 4.356 ± 0.121 | 4.208 ± 0.089 | 4.272 ± 0.068 | 4.266 ± 0.101 | 4.195 ± 0.053 |
| AGILE-COMET (layer=6) | 3.772 ± 0.053 | 3.795 ± 0.085 | 3.672 ± 0.052 | 3.651 ± 0.039 | 3.681 ± 0.049 |
| AGILE-COMET (layer=8) | 3.856 ± 0.078 | 3.766 ± 0.053 | 3.842 ± 0.062 | 3.815 ± 0.099 | 3.811 ± 0.060 |
| AGILE-COMET (layer=10) | 3.860 ± 0.092 | 3.806 ± 0.101 | 3.814 ± 0.050 | 3.815 ± 0.099 | 3.845 ± 0.068 |
| STRATA (ours) | 4.740 ± 0.141 | 4.666 ± 0.075 | 4.902 ± 0.117 | 5.118 ± 0.083 | 5.518 ± 0.234 |
| *LC2024 (N2a)* |  |  |  |  |  |
| AGILE-Metaddata | 0.580 ± 0.026 | 0.539 ± 0.012 | 0.581 ± 0.039 | 0.557 ± 0.021 | 0.624 ± 0.047 |
| AGILE-MLP | 1.700 ± 0.045 | 1.650 ± 0.047 | 1.651 ± 0.028 | 1.622 ± 0.059 | 1.648 ± 0.033 |
| AGILE-COMET (layer=6) | 1.478 ± 0.065 | 1.441 ± 0.071 | 1.465 ± 0.077 | 1.450 ± 0.056 | 1.424 ± 0.038 |
| AGILE-COMET (layer=8) | 1.457 ± 0.045 | 1.421 ± 0.035 | 1.467 ± 0.058 | 1.453 ± 0.035 | 1.466 ± 0.024 |
| AGILE-COMET (layer=10) | 1.456 ± 0.026 | 1.526 ± 0.042 | 1.518 ± 0.062 | 1.524 ± 0.030 | 1.484 ± 0.042 |
| STRATA (ours) | 1.973 ± 0.048 | 2.053 ± 0.067 | 2.060 ± 0.084 | 2.158 ± 0.121 | 2.255 ± 0.125 |

**Table 10:** Performance of STRATA with different positional encodings.

| Metric<br>dataset | MSE (↓) | | | | PCC (↑) | | | |
| --- | --- | --- | --- | --- | --- | --- | --- | --- |
|  | A549 | YZ2022 | LC2024 (N2a) | LC2024 (HEK293) | A549 | YZ2022 | LC2024 (N2a) | LC2024 (HEK293) |
| random split |  |  |  |  |  |  |  |  |
| random | 0.7197 ± 0.0424 | 0.8647 ± 0.1455 | 3.4426 ± 0.4141 | 4.7942 ± 0.2014 | 0.6796 ± 0.0282 | 0.3670 ± 0.0495 | 0.2886 ± 0.0646 | 0.2428 ± 0.0384 |
| APE | 0.5503 ± 0.0200 | 0.4624 ± 0.0620 | 1.1075 ± 0.2099 | 2.6291 ± 0.5830 | 0.7767 ± 0.0157 | 0.7469 ± 0.0587 | 0.8487 ± 0.0289 | 0.7132 ± 0.0642 |
| RoPE | 0.5511 ± 0.0420 | 0.4927 ± 0.0983 | 1.5120 ± 0.2974 | 2.9154 ± 0.4961 | 0.7711 ± 0.0232 | 0.7290 ± 0.0576 | 0.7942 ± 0.0462 | 0.6892 ± 0.0606 |
| GBF (comet) | 0.5778 ± 0.0279 | 0.4817 ± 0.1165 | 1.1939 ± 0.1555 | 2.8338 ± 0.4584 | 0.7606 ± 0.0189 | 0.7280 ± 0.0703 | 0.8325 ± 0.0319 | 0.6832 ± 0.0531 |
| RiPE (ours) | 0.4873 ± 0.0290 | 0.3698 ± 0.0667 | 0.7953 ± 0.1295 | 2.5155 ± 0.4417 | 0.8030 ± 0.0128 | 0.8015 ± 0.0452 | 0.8863 ± 0.0225 | 0.7269 ± 0.0558 |
| gain | 11.4483% | 20.0260% | 28.1896% | 4.3209% | 3.3861% | 7.3102% | 4.4303% | 1.9209% |
| unseen molecule split |  |  |  |  |  |  |  |  |
| random | 1.0007 ± 0.0877 | 1.0731 ± 0.6086 | 3.7470 ± 1.2825 | 6.7951 ± 1.3772 | 0.5504 ± 0.0576 | -0.0538 ± 0.1673 | -0.0418 ± 0.1014 | 0.0593 ± 0.1040 |
| APE | 0.6947 ± 0.1012 | 0.9871 ± 0.4978 | 2.9962 ± 1.7400 | 5.5448 ± 2.2166 | 0.7292 ± 0.0241 | 0.2741 ± 0.1713 | 0.4863 ± 0.1461 | 0.4115 ± 0.2084 |
| RoPE | 0.6917 ± 0.0866 | 1.0367 ± 0.5597 | 3.8003 ± 1.5614 | 5.1754 ± 1.5710 | 0.7252 ± 0.0248 | 0.2559 ± 0.2267 | 0.4153 ± 0.2369 | 0.4553 ± 0.1534 |
| GBF (comet) | 0.7543 ± 0.1319 | 1.0688 ± 0.5353 | 3.2946 ± 1.5187 | 6.3435 ± 1.7618 | 0.6977 ± 0.0369 | 0.0543 ± 0.1936 | 0.3902 ± 0.2189 | 0.3409 ± 0.1429 |
| RiPE (ours) | 0.6558 ± 0.0966 | 0.8652 ± 0.3153 | 2.8728 ± 1.1652 | 4.5784 ± 0.8397 | 0.7415 ± 0.0326 | 0.3451 ± 0.2211 | 0.5051 ± 0.1682 | 0.5209 ± 0.0977 |
| gain | 5.1901% | 12.3493% | 4.1186% | 11.5353% | 1.6868% | 25.9030% | 3.8659% | 14.4081% |
| unseen IL ratio split |  |  |  |  |  |  |  |  |
| random | 0.6539 ± 0.0573 | 1.1240 ± 0.5371 | 4.2737 ± 1.4860 | 5.0135 ± 1.0587 | 0.7005 ± 0.0305 | 0.3564 ± 0.1680 | 0.2666 ± 0.2224 | 0.1744 ± 0.2426 |
| APE | 0.7762 ± 0.1465 | 0.9678 ± 0.4459 | 2.6219 ± 1.6446 | 5.3475 ± 2.4358 | 0.6475 ± 0.0868 | 0.4825 ± 0.1294 | 0.6469 ± 0.1393 | 0.3988 ± 0.1498 |
| RoPE | 0.9527 ± 0.1718 | 0.9229 ± 0.3500 | 2.3285 ± 0.7900 | 4.6465 ± 1.0279 | 0.5189 ± 0.1318 | 0.5362 ± 0.1622 | 0.6430 ± 0.1645 | 0.3561 ± 0.1288 |
| GBF (comet) | 0.7902 ± 0.1710 | 1.0093 ± 0.4835 | 2.6038 ± 1.2074 | 4.3791 ± 1.1059 | 0.6416 ± 0.0836 | 0.4757 ± 0.1578 | 0.6389 ± 0.1729 | 0.4236 ± 0.1050 |
| RiPE (ours) | 0.6391 ± 0.0909 | 0.5487 ± 0.2989 | 1.5142 ± 0.5878 | 3.5891 ± 0.8104 | 0.7236 ± 0.0837 | 0.7142 ± 0.1136 | 0.7636 ± 0.1317 | 0.5392 ± 0.1484 |
| gain | 2.2633% | 40.5461% | 34.9710% | 18.0402% | 3.2976% | 33.1966% | 18.0399% | 27.2899% |
| unseen HL ratio split |  |  |  |  |  |  |  |  |
| random | 0.7940 ± 0.1537 | 0.7354 ± 0.3494 | 2.7914 ± 2.9455 | 4.8637 ± 0.7120 | 0.6583 ± 0.0527 | 0.2701 ± 0.1143 | 0.0691 ± 0.2070 | 0.1704 ± 0.0731 |
| APE | 0.8649 ± 0.1554 | 0.6075 ± 0.3380 | 2.3630 ± 3.4012 | 3.8273 ± 1.9342 | 0.6324 ± 0.0715 | 0.4580 ± 0.2005 | 0.5106 ± 0.3182 | 0.5479 ± 0.1363 |
| RoPE | 1.0746 ± 0.2768 | 0.5743 ± 0.2319 | 2.1462 ± 3.4035 | 3.4615 ± 1.0842 | 0.5451 ± 0.1150 | 0.4769 ± 0.0947 | 0.5434 ± 0.2845 | 0.5885 ± 0.0739 |
| GBF (comet) | 0.8627 ± 0.1015 | 0.8774 ± 0.3912 | 2.3974 ± 2.2657 | 4.4027 ± 1.7842 | 0.6443 ± 0.0597 | 0.3758 ± 0.1469 | 0.4541 ± 0.2453 | 0.4965 ± 0.1178 |
| RiPE (ours) | 0.6807 ± 0.0461 | 0.4757 ± 0.1957 | 1.6280 ± 1.9178 | 3.2386 ± 0.8974 | 0.7288 ± 0.0332 | 0.5404 ± 0.0801 | 0.5963 ± 0.1698 | 0.5983 ± 0.0906 |
| gain | 14.2695% | 17.1687% | 24.1450% | 6.4394% | 10.7094% | 13.3152% | 9.7350% | 1.6653% |
| unseen CHOL ratio split |  |  |  |  |  |  |  |  |
| random | 0.7546 ± 0.0878 | 0.9643 ± 0.1761 | 3.4237 ± 0.4254 | 5.1925 ± 1.0435 | 0.6436 ± 0.0454 | 0.3126 ± 0.0797 | 0.3054 ± 0.0668 | 0.2224 ± 0.0782 |
| APE | 0.8464 ± 0.1862 | 0.8020 ± 0.1969 | 2.2795 ± 1.1699 | 5.1479 ± 1.7936 | 0.6457 ± 0.0472 | 0.5518 ± 0.2878 | 0.6420 ± 0.2636 | 0.3903 ± 0.2010 |
| RoPE | 1.0389 ± 0.1793 | 0.7641 ± 0.1655 | 2.5841 ± 0.5947 | 4.4303 ± 1.2691 | 0.5282 ± 0.0608 | 0.5918 ± 0.0933 | 0.6183 ± 0.0984 | 0.4704 ± 0.0680 |
| GBF (comet) | 0.8659 ± 0.1832 | 0.7104 ± 0.1693 | 2.0860 ± 1.1002 | 4.4235 ± 1.7884 | 0.6161 ± 0.0667 | 0.6246 ± 0.0956 | 0.7261 ± 0.1311 | 0.4862 ± 0.1666 |
| RiPE (ours) | 0.6290 ± 0.0987 | 0.4084 ± 0.0934 | 1.3402 ± 0.4061 | 3.4368 ± 1.2848 | 0.7245 ± 0.0435 | 0.7965 ± 0.0498 | 0.8258 ± 0.0557 | 0.5823 ± 0.1571 |
| gain | 16.6446% | 42.5113% | 35.7526% | 22.3059% | 12.2038% | 27.5216% | 13.7309% | 19.7655% |
| unseen PEG ratio split |  |  |  |  |  |  |  |  |
| random | 0.8966 ± 0.1601 | 1.0658 ± 0.2663 | 3.4770 ± 0.9770 | 4.5788 ± 1.6576 | 0.6384 ± 0.0265 | 0.3324 ± 0.0387 | 0.3587 ± 0.0751 | 0.2111 ± 0.1300 |
| APE | 0.9390 ± 0.1406 | 0.7364 ± 0.2695 | 1.6517 ± 0.7810 | 4.4712 ± 1.8518 | 0.5655 ± 0.1040 | 0.6626 ± 0.0788 | 0.7155 ± 0.1346 | 0.3298 ± 0.2034 |
| RoPE | 1.1835 ± 0.2070 | 0.7787 ± 0.2455 | 1.4498 ± 0.5326 | 4.2721 ± 1.4487 | 0.4488 ± 0.1335 | 0.6297 ± 0.0652 | 0.7195 ± 0.1305 | 0.3490 ± 0.1972 |
| GBF (comet) | 0.8605 ± 0.1417 | 0.7526 ± 0.2317 | 1.1345 ± 0.5881 | 4.9144 ± 2.5263 | 0.5723 ± 0.0901 | 0.6303 ± 0.0953 | 0.6270 ± 0.1182 | 0.3352 ± 0.1512 |
| RiPE (ours) | 0.6694 ± 0.1668 | 0.5660 ± 0.0937 | 1.1286 ± 0.1761 | 3.9178 ± 0.9480 | 0.7121 ± 0.0481 | 0.7441 ± 0.0767 | 0.8069 ± 0.0303 | 0.3551 ± 0.2069 |
| gain | 22.2080% | 23.1396% | 22.1548% | 8.2933% | 11.5445% | 12.3000% | 12.1473% | 1.7479% |
| unseen IL to cargo ratio split |  |  |  |  |  |  |  |  |
| random | 1.5642 ± 0.2355 | 0.8604 ± 0.0484 | 3.1735 ± 0.5496 | 5.1925 ± 1.0435 | 0.2856 ± 0.0456 | 0.4115 ± 0.0424 | 0.3034 ± 0.0136 | 0.2224 ± 0.0782 |
| APE | 1.4289 ± 0.2903 | 0.6178 ± 0.1104 | 1.8158 ± 0.6145 | 5.1479 ± 1.7936 | 0.3032 ± 0.2078 | 0.6714 ± 0.0928 | 0.7423 ± 0.0945 | 0.3903 ± 0.2010 |
| RoPE | 1.4026 ± 0.1974 | 0.6652 ± 0.1555 | 2.8879 ± 0.8928 | 4.4303 ± 1.2691 | 0.3647 ± 0.2234 | 0.6282 ± 0.1112 | 0.5289 ± 0.1917 | 0.4704 ± 0.0680 |
| GBF (comet) | 1.3920 ± 0.1619 | 0.8911 ± 0.1953 | 2.0034 ± 0.4053 | 4.6761 ± 1.2118 | 0.2931 ± 0.2395 | 0.4992 ± 0.1220 | 0.7056 ± 0.0539 | 0.5200 ± 0.1014 |
| RiPE (ours) | 1.3575 ± 0.2834 | 0.4735 ± 0.1012 | 1.3321 ± 0.2225 | 3.9980 ± 1.3497 | 0.4910 ± 0.1011 | 0.7524 ± 0.0395 | 0.8084 ± 0.0402 | 0.6120 ± 0.1040 |
| gain | 2.4784% | 23.3571% | 26.6384% | 9.7578% | 34.6312% | 12.0643% | 8.9048% | 17.6923% |

##### Per-method analysis

AGILE-Metadata is the fastest among all methods because it routes only the ionizable lipid through the molecule encoder, while the remaining three lipid components are represented as metadata, which substantially reduces encoder forward passes. AGILE-COMET passes all four molecular components through the molecule encoder, but the encoder parameters are frozen during training, which keeps the cost low; we observe a mild upward trend in epoch time as the number of transformer layers increases, although this effect is minor due to the lightweight transformer and the short input sequence lengths. AGILE-MLP, despite its structurally simpler aggregation head, incurs a noticeably higher cost because it encodes all four molecules with a fully learnable molecule encoder, which dominates the per-epoch training time. STRATA encodes all four molecules through the molecule encoder twice, once with frozen parameters and once with learnable parameters, yet its training time is comparable to that of AGILE-MLP, since only one of the two encoder passes contributes gradients, and its transformer module remains lightweight with few layers and short sequence lengths.

##### Overall comparison

Across all configurations, STRATA exhibits the longest per-epoch training time and is more sensitive to the hidden dimension size than the other methods, which is consistent with its dual-encoder design. However, the gap between STRATA and the strongest baseline in terms of speed, AGILE-MLP, remains small, indicating that the additional encoder pass and the alignment objective introduce only a modest computational overhead. Considering the consistent and substantial accuracy improvements that STRATA delivers over all baselines, we view this overhead as a favorable trade-off between computational cost and predictive performance.

#### C.4 Full results for Positional Encodings

We report the results on the additional splits that were omitted from Section 4.3 due to space constraints. As previously discussed, RiPE consistently improves performance not only in the in-distribution setting but also under the unseen molecule and unseen ratio splits, demonstrating that its benefits transfer broadly across distribution-shift scenarios. Although not universal, the gains under the unseen ratio splits tend to be larger than those observed on the random split, which we attribute to RiPE’s ability to encode the inherent nature of ratios as an inductive bias, thereby aiding generalization along the ratio axis.

#### C.5 Cross-dataset Experiment

##### Experimental setup and motivation

We conduct a cross-dataset evaluation in which all components of STRATA except the prediction head are trained on a single source dataset under the random split, and the prediction head is subsequently finetuned on each target dataset. The diagonal entries of Table 11, shaded in gray, correspond to the case in which the source and target datasets are identical and the entire model is fully trained on that dataset; these results are therefore identical to those reported in the main experiment (Table 1). This experiment is more challenging than the cross-cell-type experiment in Section 4.5: in the latter, the data source is held fixed and only the cell type is varied, so external factors affecting the LNP itself, such as the formulation protocol, the range of compositional ratios, and the set of molecular species, are effectively controlled. The cross-cell-type experiment thus measures the extent to which the learned representation captures cell-type-agnostic, transferable properties of LNP formulations. In contrast, the cross-dataset experiment introduces substantial shifts in the formulation protocol, the supported ranges of compositional ratios, and the molecular libraries themselves, and therefore probes whether STRATA can still produce useful representations under markedly more demanding conditions.

**Table 11:** For the datasets in our main experiment, evalutate performance when only finetuning prediction heads.

| Metric<br>dataset | MSE (↓) | | | | PCC (↑) | | | |
| --- | --- | --- | --- | --- | --- | --- | --- | --- |
|  | A549 | YZ2022 | LC2024 (N2a) | LC2024 (HEK293) | A549 | YZ2022 | LC2024 (N2a) | LC2024 (HEK293) |
| Trained with A549 |  |  |  |  |  |  |  |  |
| AGILE-Metadata | 0.5644 ± 0.0263 | 0.8043 ± 0.1238 | 3.1837 ± 0.3438 | 5.4873 ± 0.3834 | 0.7610 ± 0.0180 | 0.4473 ± 0.0576 | 0.4742 ± 0.0436 | 0.1842 ± 0.0534 |
| AGILE-MLP | 0.5325 ± 0.0399 | 0.7877 ± 0.1378 | 2.8943 ± 0.3582 | 5.2613 ± 0.3587 | 0.7792 ± 0.0139 | 0.4607 ± 0.0548 | 0.4914 ± 0.0429 | 0.1601 ± 0.0702 |
| AGILE-COMET | 1.0022 ± 0.0657 | 0.9780 ± 0.1624 | 3.6385 ± 0.4771 | 4.6424 ± 0.3433 | 0.5019 ± 0.0340 | 0.1469 ± 0.0560 | 0.1703 ± 0.0562 | 0.2903 ± 0.0465 |
| STRATA (ours) | 0.4873 ± 0.0290 | 0.7766 ± 0.1065 | 1.9804 ± 0.3038 | 3.8437 ± 0.2424 | 0.8030 ± 0.0128 | 0.4781 ± 0.0416 | 0.6825 ± 0.0635 | 0.4918 ± 0.0323 |
| Trained with YZ2022 |  |  |  |  |  |  |  |  |
| AGILE-Metadata | 0.8086 ± 0.0376 | 0.6908 ± 0.0783 | 1.7687 ± 0.2367 | 3.6539 ± 0.1635 | 0.6368 ± 0.0241 | 0.5500 ± 0.0729 | 0.7470 ± 0.0521 | 0.5737 ± 0.0483 |
| AGILE-MLP | 0.7811 ± 0.0358 | 0.4007 ± 0.0830 | 1.6542 ± 0.2580 | 3.2729 ± 0.1975 | 0.6482 ± 0.0247 | 0.7748 ± 0.0460 | 0.7550 ± 0.0503 | 0.5988 ± 0.0471 |
| AGILE-COMET | 1.2597 ± 0.0746 | 0.5127 ± 0.0466 | 1.9088 ± 0.1984 | 4.2034 ± 0.3249 | 0.2888 ± 0.0590 | 0.7040 ± 0.0507 | 0.7018 ± 0.0514 | 0.4125 ± 0.0378 |
| STRATA (ours) | 0.8684 ± 0.0506 | 0.3698 ± 0.0667 | 1.5568 ± 0.3139 | 3.0842 ± 0.1962 | 0.5938 ± 0.0228 | 0.8015 ± 0.0452 | 0.7659 ± 0.0508 | 0.6255 ± 0.0335 |
| Trained with LC2024 (N2a) |  |  |  |  |  |  |  |  |
| AGILE-Metadata | 0.9116 ± 0.0542 | 0.4766 ± 0.0838 | 2.1385 ± 0.3542 | 3.9523 ± 0.3788 | 0.5573 ± 0.0344 | 0.7138 ± 0.0434 | 0.6660 ± 0.0698 | 0.4773 ± 0.0637 |
| AGILE-MLP | 0.8726 ± 0.0462 | 0.4504 ± 0.0921 | 1.5726 ± 0.3292 | 3.8451 ± 0.4082 | 0.5913 ± 0.0346 | 0.7393 ± 0.0475 | 0.7683 ± 0.0577 | 0.4909 ± 0.0741 |
| AGILE-COMET | 1.1967 ± 0.0812 | 0.5313 ± 0.0538 | 1.5963 ± 0.2253 | 4.2320 ± 0.3175 | 0.3236 ± 0.0447 | 0.6811 ± 0.0627 | 0.7595 ± 0.0426 | 0.4040 ± 0.0412 |
| STRATA (ours) | 0.8074 ± 0.0555 | 0.4317 ± 0.0774 | 0.7953 ± 0.1295 | 3.0180 ± 0.2486 | 0.6330 ± 0.0183 | 0.7532 ± 0.0535 | 0.8863 ± 0.0225 | 0.6400 ± 0.0360 |
| Trained with LC2024 (HEK293) |  |  |  |  |  |  |  |  |
| AGILE-Metadata | 0.8859 ± 0.0461 | 0.9004 ± 0.1842 | 2.7428 ± 0.3376 | 3.8305 ± 0.2649 | 0.5729 ± 0.0388 | 0.3233 ± 0.0442 | 0.4872 ± 0.0653 | 0.4884 ± 0.0528 |
| AGILE-MLP | 0.8524 ± 0.0375 | 0.8802 ± 0.1575 | 2.6853 ± 0.4117 | 3.7015 ± 0.2307 | 0.6019 ± 0.0249 | 0.3443 ± 0.0821 | 0.5384 ± 0.0625 | 0.5176 ± 0.0460 |
| AGILE-COMET | 1.3335 ± 0.0741 | 0.8907 ± 0.1375 | 2.9284 ± 0.3890 | 3.3464 ± 0.2654 | 0.0217 ± 0.0365 | 0.3256 ± 0.0298 | 0.4603 ± 0.0880 | 0.5814 ± 0.0535 |
| STRATA (ours) | 0.7631 ± 0.0389 | 0.4253 ± 0.0701 | 1.4281 ± 0.2137 | 2.5155 ± 0.4417 | 0.6565 ± 0.0192 | 0.7635 ± 0.0423 | 0.7866 ± 0.0389 | 0.7269 ± 0.0558 |

##### Cross-dataset transfer performance

As expected, the cross-dataset transfer setting yields performance that is generally below the gray-shaded full-training results, and STRATA is not the top-performing model in every individual cell of Table 11. Nevertheless, STRATA achieves the best performance in the largest number of source–target combinations, and the PCC values it attains are often surprisingly high in absolute terms. Since PCC reflects the model’s ability to recover the relative ordering of samples, these high PCC values indicate that STRATA captures the underlying tendencies of LNP behavior well, even when the absolute scale of the target distribution differs from the source. We highlight two particularly noteworthy cases. When trained on LC2024 (HEK293) and finetuned on LC2024 (N2a), STRATA achieves better MSE and PCC than every baseline that is fully trained directly on LC2024 (N2a). Similarly, when trained on LC2024 (HEK293) and finetuned on YZ2022, STRATA’s performance is on par with the strongest baseline that is fully trained on YZ2022. These observations strongly suggest that STRATA’s interaction modeling is highly effective: even when the model has never been fully trained on the target data, the representations transferred from a different dataset are sufficient to match or surpass baselines that enjoy full access to the target data.

##### Embedding analysis via Moran’s I

To further investigate why STRATA transfers so effectively across datasets, we apply t-SNE to the embeddings produced by a model that is trained on LC2024 (N2a) and then evaluated, without any finetuning, on the A549 dataset. We color each point by one of the five compositional ratios (IL, HL, CHOL, PEG, and cargo), and additionally report Moran’s I, computed on a row-normalized *k*-nearest-neighbor graph (*k* = 10) constructed in the t-SNE space, which quantifies the degree to which neighboring points share similar values of the colored attribute. As shown in Figure 6, both the qualitative t-SNE structure and the quantitative Moran’s I values demonstrate that STRATA’s sensitivity to compositional ratios is preserved when the embeddings are transferred to a different dataset, with substantially higher Moran’s I than AGILE-COMET on four of the five ratios. Taken together, these results indicate that STRATA’s representations retain their ratio-aware geometric structure even on out-of-distribution data, which we believe is a major reason for its strong cross-dataset transfer performance.

**Figure 6:**
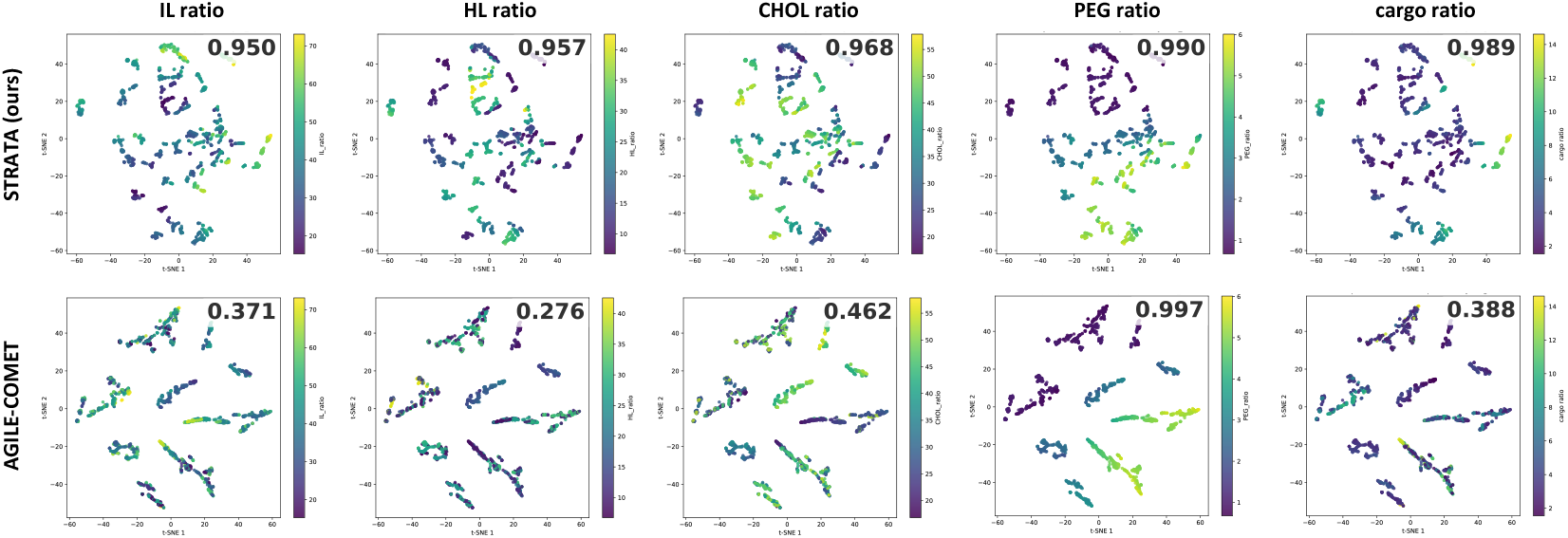
t-SNE visualization of LNP embeddings. The first row shows results from our model, STRATA, and the second row shows results from AGILE-COMET. Each column corresponds to results labeled by five different ratios. The value represents Moran’s I.

#### C.6 Baseline Comparison (respecting their own molecule encoders)

In our main experiments, we replaced the molecule encoders of all baselines with the AGILE molecule encoder in order to control for the influence of the molecule encoder itself and the amount of lipid-specific additional pretraining, because our research focuses on how to effectively integrate compositional ratios with molecular representations rather than on improving the molecule encoder. For completeness, however, we anticipate that readers may also be interested in the performance of the existing models when each baseline retains its originally proposed molecule encoder, and we therefore report these results in Table 12.

**Table 12:**
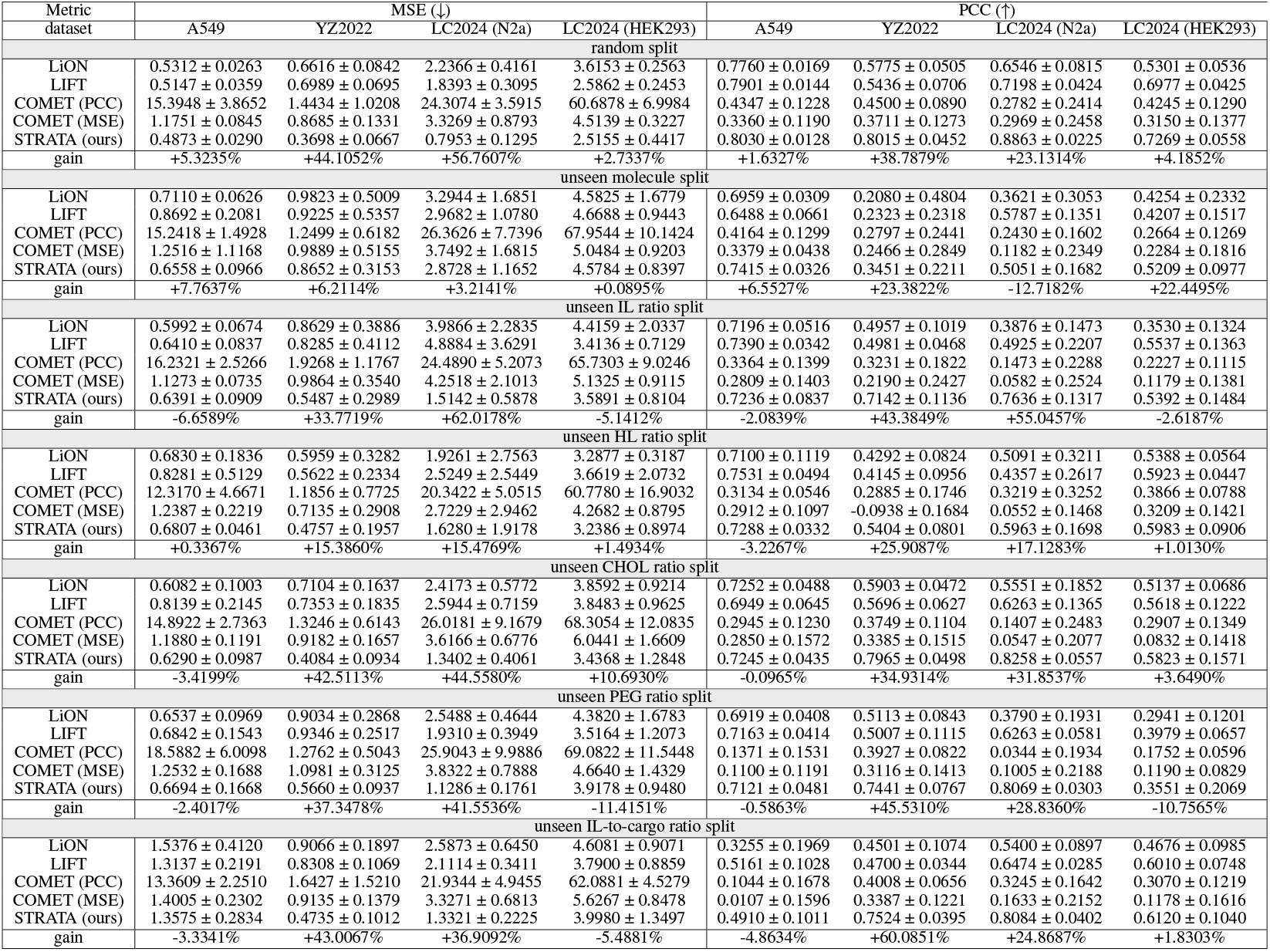
Performance of existing models, respecting their own molecule encoders.

##### Result analysis

We attribute the variation in performance across baselines to several factors, including differences in the architecture and modalities of each molecule encoder and differences in the amount of lipid-specific data used for additional pretraining of these encoders. Although STRATA is outperformed by some baselines in a few individual cases, the gain rows in Table 12 show that the corresponding gaps are generally small. In contrast, STRATA achieves wins by substantially larger performance margins in many other cases, particularly under the unseen ratio splits, which are precisely the settings that most directly stress the model’s ability to generalize along compositional axes. These observations indicate that, even when baselines are allowed to leverage their own carefully designed and separately pretrained molecule encoders, STRATA’s interaction modeling remains highly competitive overall and exhibits a clear advantage in the most challenging compositional generalization scenarios.

#### C.7 Spearman’s rank correlation coefficient

In LNP high-throughput screening, the practical objective is to identify formulations with high transfection efficiency, making sample-ranking metrics particularly relevant for evaluation. Accordingly, we report Spearman’s rank correlation coefficients in Table 13. Overall, the results show trends consistent with those observed in Section 4.2.

**Table 13:**
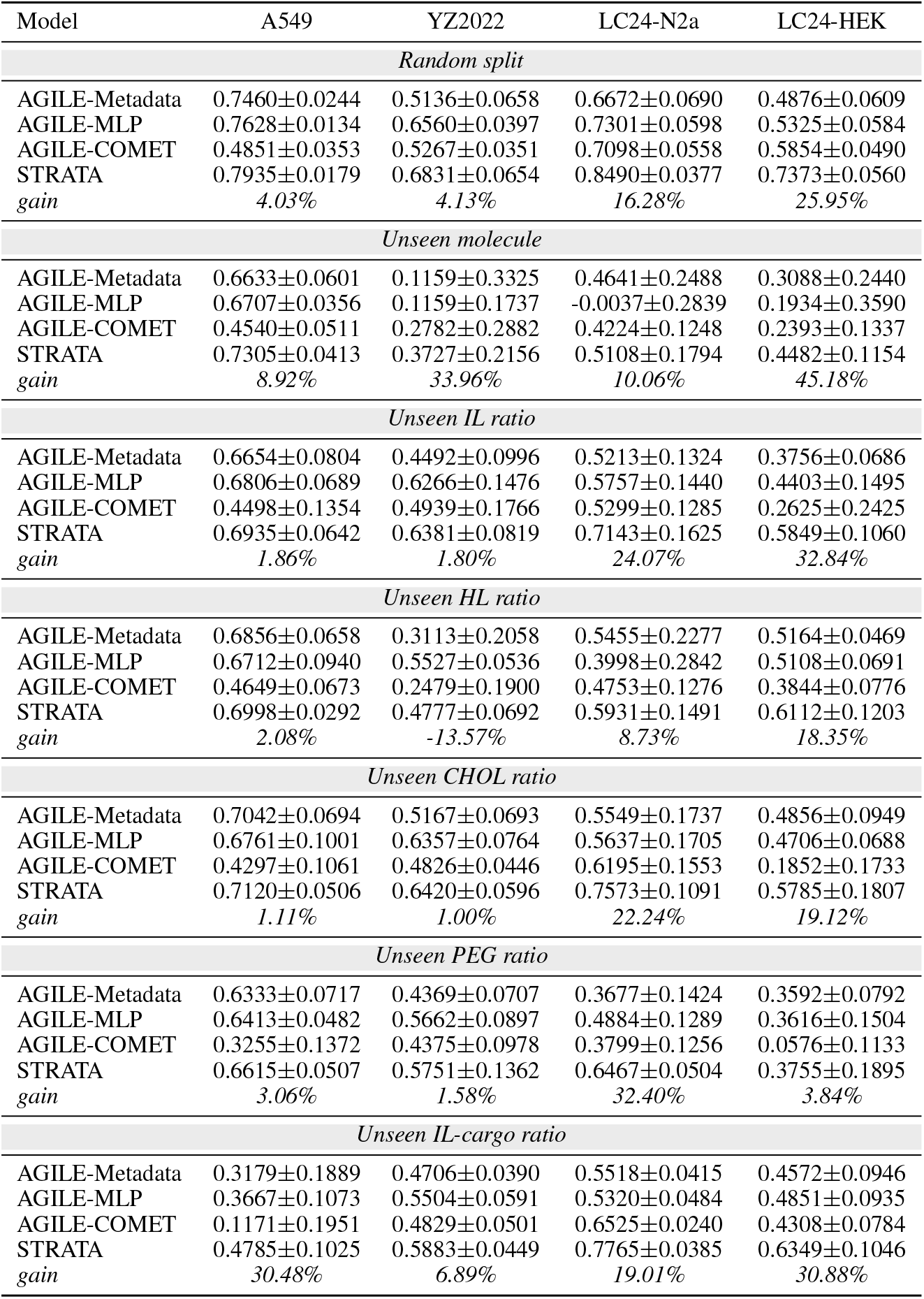
Results across dataset splits, with gain indicating STRATA’s relative improvement over the best baseline.

| Model | A549 | YZ2022 | LC24-N2a | LC24-HEK |
| --- | --- | --- | --- | --- |
| *Random split* |  |  |  |  |
| AGILE-Metadata | 0.7460±0.0244 | 0.5136±0.0658 | 0.6672±0.0690 | 0.4876±0.0609 |
| AGILE-MLP | 0.7628±0.0134 | 0.6560±0.0397 | 0.7301±0.0598 | 0.5325±0.0584 |
| AGILE-COMET | 0.4851±0.0353 | 0.5267±0.0351 | 0.7098±0.0558 | 0.5854±0.0490 |
| STRATA | 0.7935±0.0179 | 0.6831±0.0654 | 0.8490±0.0377 | 0.7373±0.0560 |
| *gain* | 4.03% | 4.13% | 16.28% | 25.95% |
| *Unseen molecule* |  |  |  |  |
| AGILE-Metadata | 0.6633±0.0601 | 0.1159±0.3325 | 0.4641±0.2488 | 0.3088±0.2440 |
| AGILE-MLP | 0.6707±0.0356 | 0.1159±0.1737 | -0.0037±0.2839 | 0.1934±0.3590 |
| AGILE-COMET | 0.4540±0.0511 | 0.2782±0.2882 | 0.4224±0.1248 | 0.2393±0.1337 |
| STRATA | 0.7305±0.0413 | 0.3727±0.2156 | 0.5108±0.1794 | 0.4482±0.1154 |
| *gain* | 8.92% | 33.96% | 10.06% | 45.18% |
| *Unseen IL ratio* |  |  |  |  |
| AGILE-Metadata | 0.6654±0.0804 | 0.4492±0.0996 | 0.5213±0.1324 | 0.3756±0.0686 |
| AGILE-MLP | 0.6806±0.0689 | 0.6266±0.1476 | 0.5757±0.1440 | 0.4403±0.1495 |
| AGILE-COMET | 0.4498±0.1354 | 0.4939±0.1766 | 0.5299±0.1285 | 0.2625±0.2425 |
| STRATA | 0.6935±0.0642 | 0.6381±0.0819 | 0.7143±0.1625 | 0.5849±0.1060 |
| *gain* | 1.86% | 1.80% | 24.07% | 32.84% |
| *Unseen HL ratio* |  |  |  |  |
| AGILE-Metadata | 0.6856±0.0658 | 0.3113±0.2058 | 0.5455±0.2277 | 0.5164±0.0469 |
| AGILE-MLP | 0.6712±0.0940 | 0.5527±0.0536 | 0.3998±0.2842 | 0.5108±0.0691 |
| AGILE-COMET | 0.4649±0.0673 | 0.2479±0.1900 | 0.4753±0.1276 | 0.3844±0.0776 |
| STRATA | 0.6998±0.0292 | 0.4777±0.0692 | 0.5931±0.1491 | 0.6112±0.1203 |
| *gain* | 2.08% | -13.57% | 8.73% | 18.35% |
| *Unseen CHOL ratio* |  |  |  |  |
| AGILE-Metadata | 0.7042±0.0694 | 0.5167±0.0693 | 0.5549±0.1737 | 0.4856±0.0949 |
| AGILE-MLP | 0.6761±0.1001 | 0.6357±0.0764 | 0.5637±0.1705 | 0.4706±0.0688 |
| AGILE-COMET | 0.4297±0.1061 | 0.4826±0.0446 | 0.6195±0.1553 | 0.1852±0.1733 |
| STRATA | 0.7120±0.0506 | 0.6420±0.0596 | 0.7573±0.1091 | 0.5785±0.1807 |
| *gain* | 1.11% | 1.00% | 22.24% | 19.12% |
| *Unseen PEG ratio* |  |  |  |  |
| AGILE-Metadata | 0.6333±0.0717 | 0.4369±0.0707 | 0.3677±0.1424 | 0.3592±0.0792 |
| AGILE-MLP | 0.6413±0.0482 | 0.5662±0.0897 | 0.4884±0.1289 | 0.3616±0.1504 |
| AGILE-COMET | 0.3255±0.1372 | 0.4375±0.0978 | 0.3799±0.1256 | 0.0576±0.1133 |
| STRATA | 0.6615±0.0507 | 0.5751±0.1362 | 0.6467±0.0504 | 0.3755±0.1895 |
| *gain* | 3.06% | 1.58% | 32.40% | 3.84% |
| *Unseen IL-cargo ratio* |  |  |  |  |
| AGILE-Metadata | 0.3179±0.1889 | 0.4706±0.0390 | 0.5518±0.0415 | 0.4572±0.0946 |
| AGILE-MLP | 0.3667±0.1073 | 0.5504±0.0591 | 0.5320±0.0484 | 0.4851±0.0935 |
| AGILE-COMET | 0.1171±0.1951 | 0.4829±0.0501 | 0.6525±0.0240 | 0.4308±0.0784 |
| STRATA | 0.4785±0.1025 | 0.5883±0.0449 | 0.7765±0.0385 | 0.6349±0.1046 |
| *gain* | 30.48% | 6.89% | 19.01% | 30.88% |

### D Broader Impact

#### Broader Impact

Lipid nanoparticles (LNPs) are a critical delivery vehicle for mRNA-based therapeutics and vaccines, and the empirical screening of LNP formulations remains a major bottleneck in their development, typically requiring extensive wet-lab experimentation that is both costly and time-consuming. By providing accurate transfection efficiency predictions that generalize to unseen molecules and unseen formulation ratios, STRATA has the potential to substantially reduce the experimental burden in early-stage LNP design. This could accelerate the development of mRNA vaccines, gene therapies, and other nucleic acid-based treatments, ultimately broadening patient access to emerging therapeutic modalities.

Beyond the immediate application, the methodological contribution of RiPE, a positional encoding tailored to continuous compositional variables—may benefit adjacent scientific domains where ratio-structured inputs are prevalent, such as polymer formulation, alloy design, catalyst composition, and drug combination therapy. In this sense, our work contributes to the broader effort of building machine learning models that respect the inductive biases of physical and chemical systems.

We also acknowledge potential risks. First, predictions from any *in silico* model, including ours, should not be used as a substitute for experimental validation, particularly for therapeutic decision-making. Over-reliance on model predictions in safety-critical pipelines could lead to suboptimal or unsafe formulations being prioritized for downstream development. Second, although our datasets cover diverse cell lines, they remain limited in chemical and biological scope; predictions on formulations far outside the training distribution should be interpreted with caution, despite the generalization improvements demonstrated in this work. Finally, as with most molecular property prediction tools, STRATA could in principle be misused to design delivery vehicles for harmful payloads. However, we note that the bottleneck for such misuse lies primarily in payload synthesis and biological deployment rather than in formulation prediction, and we believe the benefits of openly advancing LNP design tools substantially outweigh this risk. We encourage practitioners to pair model predictions with appropriate experimental validation and to follow established biosafety and ethical guidelines.

## Notes

### Competing Interest Statement

The authors have declared no competing interest.

### Summary of Updates

Revised explanations at Section A.6. Added more experiment results at the appendix.

